# A graph-embedded topic model enables characterization of diverse pain phenotypes among UK Biobank individuals

**DOI:** 10.1101/2022.01.07.475444

**Authors:** Yuening Wang, Rodrigo Benavides, Luda Diatchenko, Audrey V. Grant, Yue Li

## Abstract

Large biobank repositories of clinical conditions and medications data open opportunities to investigate the phenotypic disease network. To enable systematic investigation of entire structured phenomes, we present graph embedded topic model (GETM). Our contributions are two folds in terms of method and applications. On the methodology side, we offer two main contributions in GETM. First, to aid topic inference, we integrate existing biomedical knowledge graph information in the form of pre-trained graph embedding into the embedded topic model. Second, leveraging deep learning techniques, we developed a variational autoencoder framework to infer patient phenotypic mixture by modeling multi-modal discrete patient medical records. In particular, for interpretability, we use a linear decoder to simultaneously infer the bi-modal distributions of the disease conditions and medications. On the application side, we applied GETM to UK Biobank (UKB) self-reported clinical phenotype data, which contains 443 self-reported medical conditions and 802 self-reported medications for 457,461 individuals. Compared to existing methods, GETM demonstrates overall superior performance in imputing missing conditions and medications. Here, we focused on characterizing pain phenotypes recorded in the questionnaire of the UKB individuals. GETM accurately predicts the status of chronic musculoskeletal (CMK) pain, chronic pain by body-site, and non-specific chronic pain using past conditions and medications. Our analyses revealed not only the known pain-related topics but also the surprising predominance of medications and conditions in the cardiovascular category among the most predictive topics across chronic pain phenotypes.

## 1 INTRODUCTION

The advent of electronic health records (EHR) has started to transform the way healthcare data are recorded and used in clinical practice and in research settings. Besides free-form clinical text, most modern healthcare centers now routinely collect structured EHR data describing comprehensive aspects of care, including diagnosis, medications, treatments, laboratory test results, and other measures. Deriving coherent phenotypes from these EHR data is crucial in downstream phenome-wide association analyses (PheWAS) and may greatly improve the power of detecting genetic associations using the genome-wide association (GWA) approach [**?**]. Besides better characterizing known phenotypes (e.g. through co-morbidities or the demographics of age-of-onset), mining phenomic data also has the potential to reveal novel combinations of diseases and other variables of potential etiological interest. This will help identify specific strata of study subjects most at risk for disease or targeted for specific drug recommendations. Despite these promises, clinical phenotype data sources remain underused [**?**]. As genotype and deep phenotype data become increasingly available through consortia or large government-funded cohorts such as the UK Biobank (UKB) [**?**] and the 100,000 Genomes Project [**?**], there is an urgent need for an automatic and accurate phenotyping tool to accelerate novel disease co-morbidity discoveries and improve the yield of GWA studies for complex phenotypes and diseases in humans.

Among many machine learning approaches, topic models [**?**] stand out as a particularly well-adapted framework for automatic phenotyping. They are extremely efficient at modeling sparse and discrete data such as text documents. Topic models were originally developed to discover patterns of word usages from corpuses of text documents by accomplishing two related tasks: (1) inferring a set of latent categorical distributions over the vocabulary (i.e., topics); (2) using these latent topic distributions to infer topic mixture memberships of each document, thereby connecting them under similar topical themes. In our context, we consider EHR as our documents and the diagnostic and medication codes as our vocabulary.

Several topic methods were developed recently for effectively mining EHR data [**?, ?**]. However, most existing topic models are unable to incorporate existing biomedical knowledge graphs, which manifest in several forms such as disease taxonomy and drug classification systems. Knowledge Graph Embedding LDA (KGE-LDA) [**?**] models the distribution of the word embedding learned from TransE [**?**] on a words-by-words relational graph. Latent-feature LDA [**?**] and Embedded Topic Model (ETM) [**?**] use the word embedding to compose the topic distribution. These methods were applied to standard benchmark corpus data and only works with one data modality as opposed to the multimodal patient electronic medical record data (e.g., disease conditions and medications). Additionally, except for ETM, the above related topic models use traditional inference algorithms (e.g., Gibbs sampling or mean-field variational inference) to infer topic distributions. Therefore, these models have limited flexibility to capture the non-linear connections between observed EHR codes and the underlying patient phenotypes.

In this paper, we present a graph-embedded topic model (GETM) for learning phenotypes from heterogeneous EHR data by leveraging biomedical graph information. As the main method contribution, GETM seamlessly integrates two existing models: ETM [**?**] and node2vec [**?**]. Briefly, we first use node2vec to learn the condition and medication embeddings based on their taxonomic information; then, we incorporate these embeddings into the ETM, which tri-factorizes the individuals-by-conditions/medications matrix into individuals-by-topics, topics-by-embedding, and embedding-by-conditions/medications matrices. The distribution of the individuals-by-topics is approximated by the output of a feedforward neural network using the amortized variational inference technique while fixing the embedding-by-conditions/medications matrices to the node2vec-learned node embedding from conditions/medications taxonomical graphs, respectively.

As a proof-of-concept, we applied GETM to UKB phenotype data, where 457,461 individuals of European descent from across the United Kingdom were deeply phenotyped through extensive self-report based questionnaires for about 443 well-defined phenotype conditions and 802 active ingredients of medications [**?**]. We then turned to a more focused analysis on predicting and characterizing different pain phenotypes. Chronic pain is the result of dysfunction of the nociceptive circuitry leading to sustained perception of pain. Chronic pain is highly prevalent in aging populations affecting up to 50% of older adults (*>*65 years old) [**?**] and decreases the overall mental and emotional health of affected individuals [**?**]. Using GETM, we provide a refreshing view of pain phenotypes, considered as CMK pain, chronic pain by body site, non-specific acute and chronic pain, and the transition from acute to chronic pain by making use of phenotype data from subsequent visits on a subset of the UKB study population. By correlating the inferred GETM-topics and pain phenotypes across the UKB subjects, we discover not only the known pain-related conditions and medications among the highly predictive topics but also novel combinations after removing labels of known pain-related conditions and analgesics.

## 2 RESULTS

### 2.1 Graph embedded topic model overview

GETM models the distribution of medications and conditions as discrete clinical features for each individual. For a given subject of study, the expected rate of each feature is determined by both the logistic-normal latent patient’s topic mixture and the point-estimate latent topic distributions over the features. The goal of GETM is to approximately infer the distributions of these latent variables. To this end, we carry out an amortized variational inference [**?**] in two steps (**Fig**. 1). In the first step, to infer the topic mixture of a given patient, we provide to a feedforward neural network (i.e., the encoder) the binary vector of the individual’s observed discrete features. We then sample the topic mixture from a variational Gaussian distribution with the mean and standard deviation computed by the encoder. In the second step, we decode the sampled topic mixture back to the original conditions and medications. We used two linear decoders each with separate topic/feature embeddings for medications and conditions, respectively. Specifically, each decoder tri-factorizes the individuals-by-features matrix respectively into individuals-by-topics (***θ***), topics-by-embeddings (***α***^(*t*)^), and embeddings-by-features (medication or condition) (***ρ***^(*t*)^) matrices (*t ∈ {med, cond}*). Notably, the two linear decoders share the patient-level topic mixture ***θ*** while ***α***^(*t*)^ and ***ρ***^(*t*)^ are learned separately. Importantly, the medication embedding ***ρ***^(*med*)^ and condition embedding ***ρ***^(*cond*)^ are pre-trained by node2vec [**?**] from the taxonomic tree-structured graphs of conditions and medications while the two sets of topic embedding counterparts (***α***^(*med*)^, ***α***^(*cond*)^) are directly learned from the UKB participant data by GETM. This tri-factorization design allows for exploring topics, study subjects and relationships among conditions and medications in a highly interpretable way.

**Figure 1:**
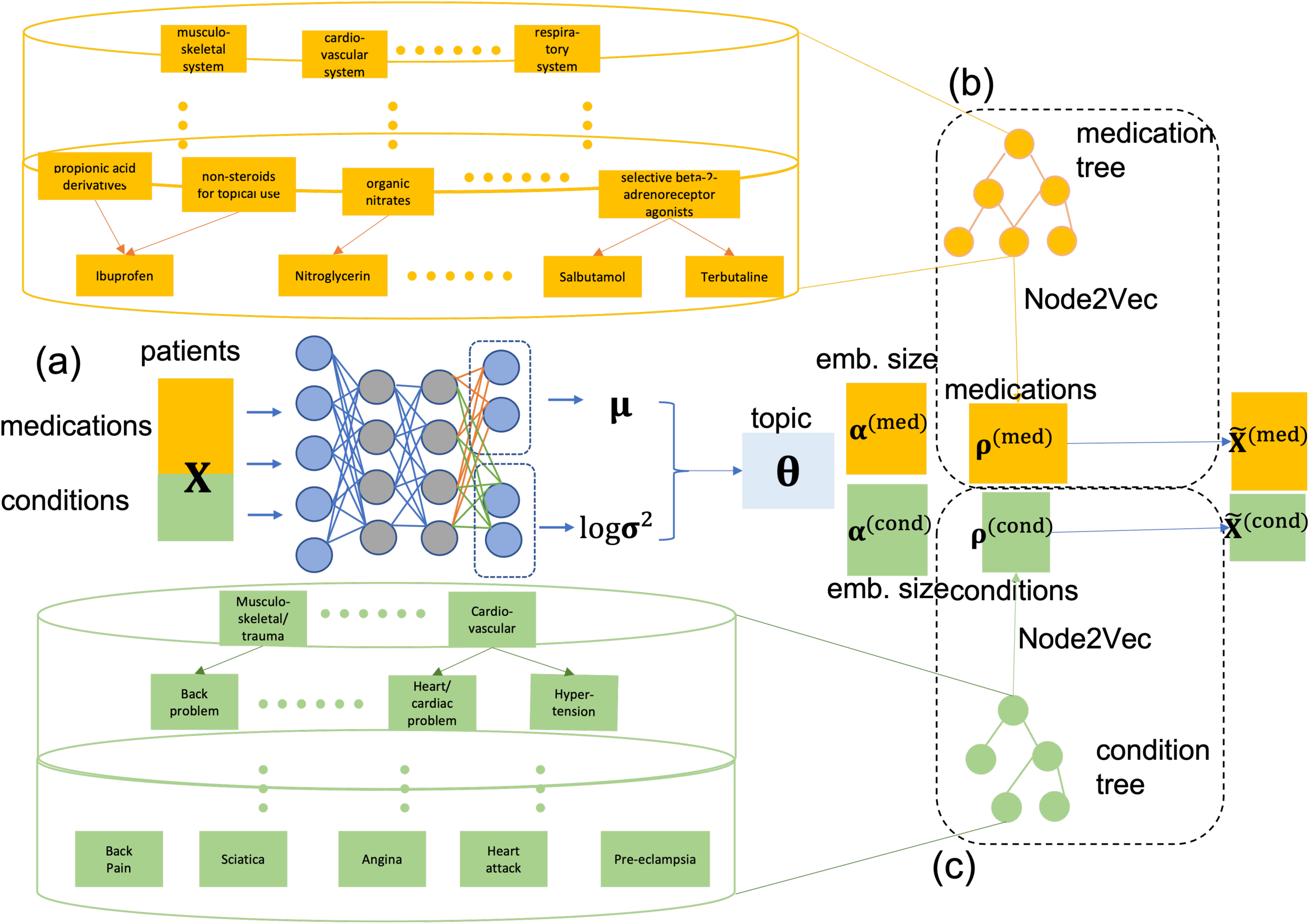
Overview of Graph-embedded topic model and its application on UKB phenotype data. The UKB data consists of 443 conditions and 802 medications for 457,461 individuals. We developed GETM to model these data although our GETM can be applied to other datasets as well. (**a**) GETM training. GETM is a variational autoencoder (VAE) model. The neural network encoder takes individuals’ condition and medication information as input and produces the variational mean *μ* and variance *σ*^2^ for the patient topic mixtures ***θ***. The decoder is linear and consists of two tri-factorizations. One learns medication-defined topic embedding ***α***^(*med*)^ and medication embedding ***ρ***^(*med*)^. The other learns condition-specific topic embedding ***α***^(*cond*)^ and the condition embedding ***ρ***^(*cond*)^. We separately pre-train (**b**) the embedding of medications ***ρ***^(*med*)^ and (**c**) the embedding of conditions ***ρ***^(*cond*)^ using node2vec [**?**] based on their structural meta-information. This is done learning the node embedding that maximizes the likelihood of the tree-structured relational graphs of conditions and medications.

### 2.2 Topic quality evaluation

Using data from the UKB on 457,461 individuals of European descent from the baseline visit, we trained GETM to obtain the topic embedding and conditions/medications embedding (i.e., ***α***^(*t*)^, ***ρ***^(*t*)^, respectively). As a qualitative exploratory analysis, we visualized these embeddings using UMAP (**Fig**. 2). For illustration purpose, we picked 5 topics representing diverse conditions and medications and observed that the top features under these topics belong to coherent categories of conditions or medications. For example, top medications atenolol, bisoprolol, metoprolol, nebivolol and carvedilol in topic 32 all belong to cardiovascular-system medications (**Fig**. 2d), while topic 32 is assigned to the cluster mainly composed of medications from the same category (**Fig**. 2b). Moreover, the top 5 conditions from topic 8 are all from the musculoskeletal/trauma category while the top 5 medications from topic 8 are all from the dermatological category. In contrast, ETM without using the Knowledge-Graph (KG)-informed embedding produced less interpretable topics each covering heterogeneous categories (**Supplementary Fig. ??**). Thus, GETM allows for the identification of related features from sparse, heterogeneous data in a data-driven manner.

**Figure 2:**
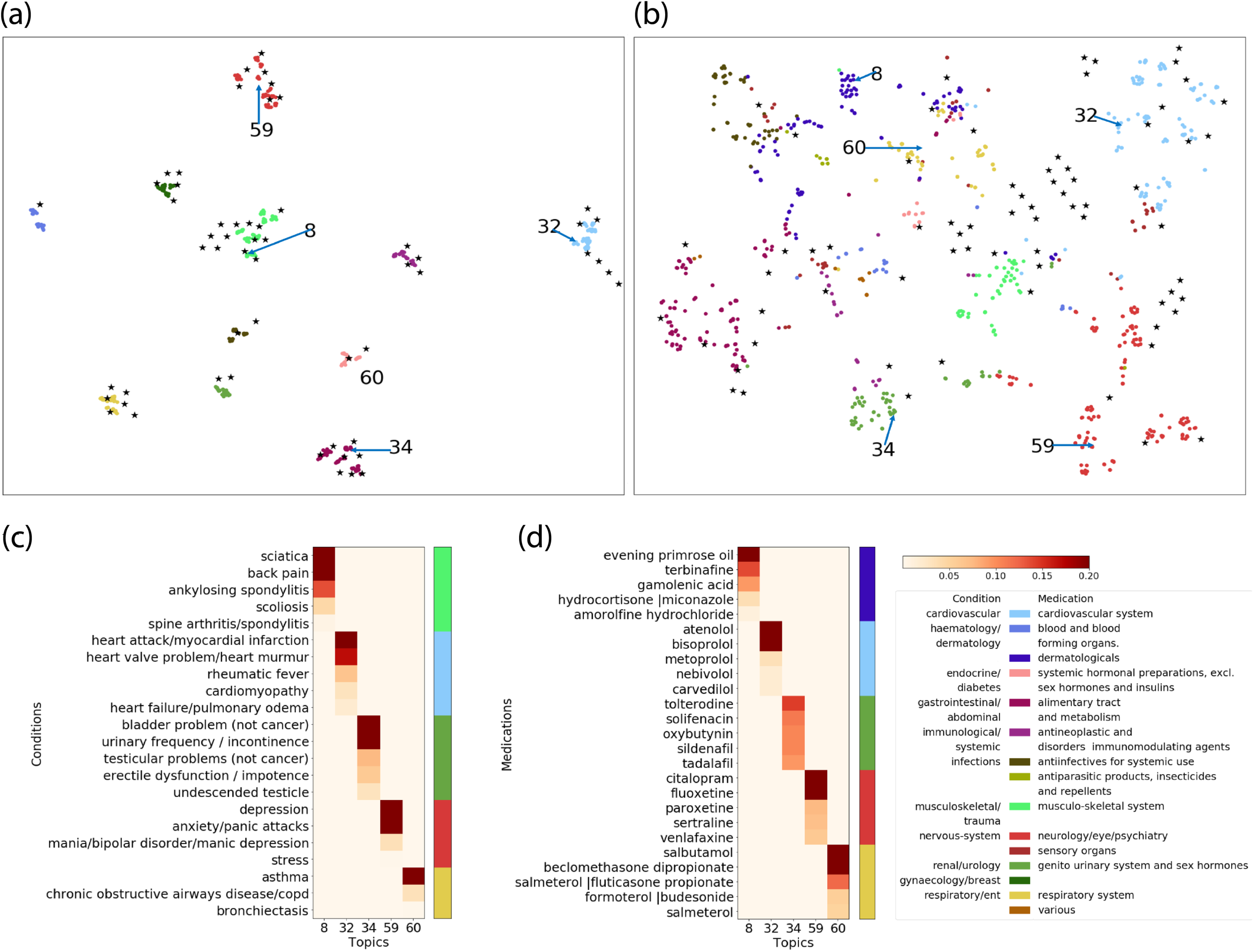
Visualizing the topic embedding and feature embedding learned by GETM on the UK Biobank data. The analysis was based on results from GETM model with both condition and medication embedding that are KG-informed using 75 topics (**Supplementary Table ??**). (**a**) Visualizing the embedding of topics and conditions. Because of the shared embedding space, we applied a single UMAP to project and visualize the condition embedding ***ρ***^(*cond*)^ and topic embedding ***α***^(*cond*)^. (**b**) Visualizing the embedding of topics and medications. Similarly, we applied a single UMAP to visualize medication embedding ***ρ***^(*med*)^ and topic embedding ***α***^(*med*)^. The solid dots on the UMAP plot are the features, and the asterisks are the topics. The points are colored by corresponding category of condition and medication or annotated by its topic number. Visualizing the UMAP in this way allows us to examine (1) the similarity among topics, (2) the similarity among features, and (3) the similarity between topics and features. As indicated by the arrows, we identified 5 topics on each UMAP and displayed their top features in panel c and d. (**c**) & (**d**) Heatmap visualization of select topics. We generated heatmaps for the top 5 conditions and the top 5 medications with the highest probability 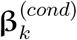 and 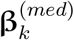 under each of the five topics. The color intensity display proportionally the probabilities. The color bars on the right indicate the categories of the conditions and medications. All four panels share the same color legend of the categories.

Given that we are leveraging embeddings of conditions and medications learned from their taxonomic graphs, we expected a higher quality of topics to be inferred by GETM compared to baseline models that do not use or use only partial graph embeddings. Topic quality was quantified as the product of topic coherence and topic diversity [**?**] (**STAR Methods**). For ease of reference, we listed the 9 models that we compared in **Supplementary Table ??**. We first evaluated the quality of the medication-defined topic (**Supplementary Table ??**). To evaluate the medication topic coherence unbiasedly, we used an external set of 59 expert-curated medication categories (**STAR Methods**) that were not part of the medication taxonomy we used in training the graph embedding for our GETM model.

We repeated our experiments 5 times each time with a different random initialization for each model and evaluated the resulting topic coherence, topic diversity, and topic quality. Overall, we observed the highest average topic quality (**Fig**. 3; **Supplementary Table ??**) for the 50-topic GETM that used the graph embeddings for both the conditions and medications (i.e., GETM in **Supplementary Table ??**). Statistically, GETM is significantly better than all of the baseline methods at p-value *<* 0.01 (one-sided t-test) except for GETM-medOnly, compared to which GETM is better at p-value *<* 0.05 (one-sided t-test). Therefore, the results suggest the benefits of using KG-informed medication embeddings in improving topic quality.

**Figure 3:**
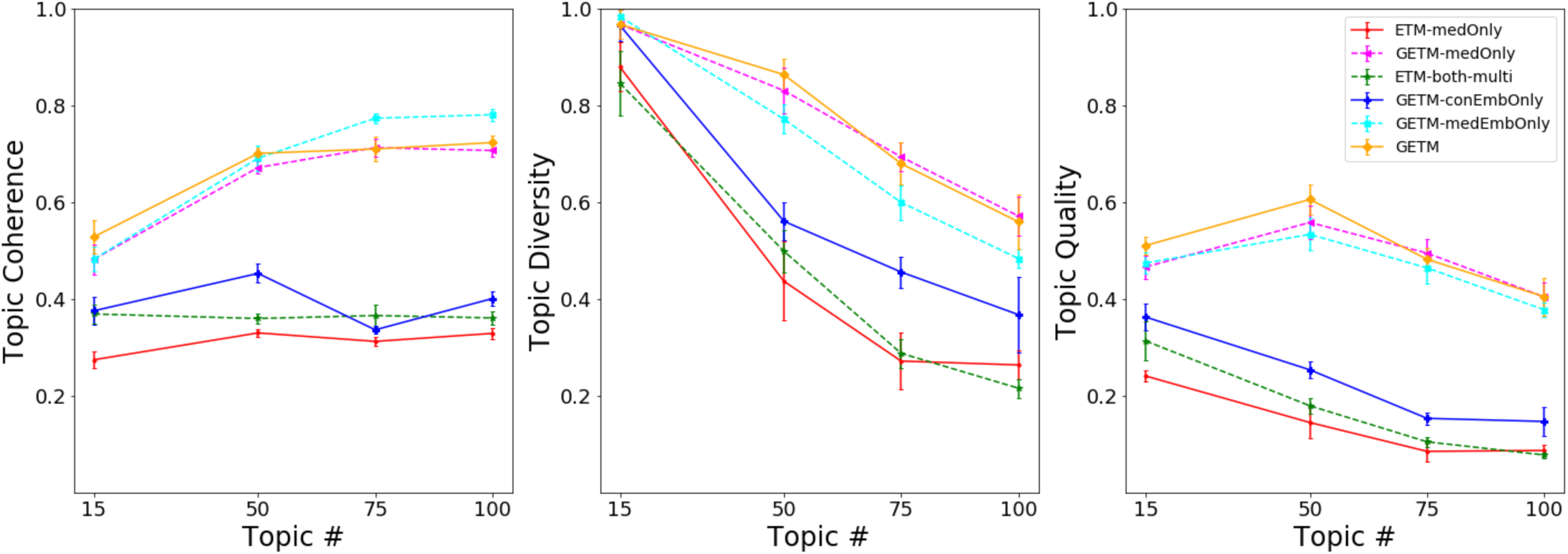
Medication-specific topic quality evaluation. We experimented with GTEM as well as 5 baseline models (**Supplementary Table ??**) with 4 predefined number of topics. To compute statistical significance between GETM and the baseline methods, we ran each model 5 times on the full UK Biobank data each with a different random initialization. The line plot displays the topic coherence, topic diversity, and topic quality, where the error bar indicates the standard deviations over the 5 experiments.

GETM-medEmbOnly confers slightly higher topic coherence but much poorer topic diversity for medications compared to GETM. This is possibly because the condition embedding directly learned from the UKB data provides complementary supports to inferring more coherent medication topics compared to the node2vec-pretrained condition embedding from the taxonomy. Note that topic coherence measures each topic separately while topic diversity measures the differences among topics. Both metrics are important. For instance, a model that generates a set of very similar topic distributions each with high topic coherent score is much less useful than a model that generates a set of diverse topic distributions each with slightly lower topic coherence score.

We evaluated condition-derived topic quality (**Supplementary Fig. ??**). Due to the lack of external condition categories, we calculated topic coherence based on whether the top conditions from each topic were present in the same patients. One caveat of this approach is that a patient rarely exhibits multiple conditions from the same category and that many conditions can be mutually exclusive within the same patient. For example, asthma and COPD as shown in our GETM topic 60 in **Fig**. 2b are rarely observed in the same patient despite the common physiological connection between them. This is reflected in the low topic coherence among the methods. Nonetheless, we found that the GETM with pretrained embeddings of both medications and conditions (**Supplementary Table ??**) dominate other models among 3 out of 4 model settings in terms of the overall topic quality scores (**Supplementary Table ??**). GETM-medEmbOnly confers much poorer topic coherence for conditions (**Supplementary Table ??**). This suggests an overall benefits of transferring the pretrained embedding for both conditions and medications to the GETM when modeling the UKB data.

### 2.3 GETM reveals known or potentially novel condition-medication relations from UKB data

We compared the total number of unique known pairs between medications and conditions that were identified by the five models (**Supplementary Table ??, Table** 1). The full GETM extracted most pairs of correlated conditions and medications illustrated by topic number 50 (161 pairs), 75 (175 pairs), and 100 (203 pairs). We examined some of the topics with high condition-medication associations based on pharmacological knowledge (**Fig**. 2b,d). For instance, in topic 32, the top medication bisoprolol is known to be used to treat the top condition heart failure under that topic. Under topic 60, the top medication salmeterol is often prescribed to treat asthma and chronic obstructive airways (COPD) [**?**], which are the top conditions under that topic. Interestingly, although the medication solifenacin in topic 59 is not known to be indicative of depression according to DrugBank, recent research shows potential for solifenacin, along with other muscarinic antagonists, in treating patients with depression through drug repurposing [**?, ?**].

### 2.4 GETM accurately imputes UKB conditions and medications

To further ascertain the overall utility of GETM, we used GETM to reconstruct randomly masked 50% of the medications or conditions of each test individual. As a baseline, we used the observed condition or medication prevalence from the training data to impute the masked feature in the test individuals. We have also experimented with the ablated models listed in **Supplementary Table ??**. Among all models, the full GETM conferred the lowest reconstruction errors for both conditions (**Table** 2) and medications (**Supplementary Table ??**) using 100 topics with *∼*5% improvement over the second best method. The improvements are largely attributed to the use of KG-informed embeddings compared to the ablated ETM model that learn these embeddings from the data.

Moreover, we also evaluated GETM in its ability to recapitulate the medication data using only the condition information by masking all medication information of the test individuals. The full GETM outperformed all other models and gave the lowest reconstruction error (14.6125), the highest precision at top 5 predicted hits (precision@5) (0.2612), and the highest recall@5 (0.5787) (**Table** 3,4). The upper bounds, which were obtained from the GETM trained on unmasked test data achieved reconstruction error of 11.4936, precision@5 of 0.4226, and recall@5 of 0.8027. Interestingly, we found out that on average around 44% of the unobserved medications from the top 10 predicted medications by GTEM in fact have a treatment effect based on condition-medication association information extracted from Comparative Toxicogenomics Database (CTD) http://ctdbase.org/ and DrugBank https://go.drugbank.com/. In particular, we took a closer look at the 3 best predicted patients and 3 worse predicted patients by GETM (**Fig**. 4). As expected, among the best predicted patients, the top 10 predicted medications by GETM contain mostly observed medications, which are known to match the subset of the conditions of these 3 patients. Although most top predicted medications were not observed for the worst 3 predicted patients, they were known to treat the observed conditions for these patients.

**Figure 4:**
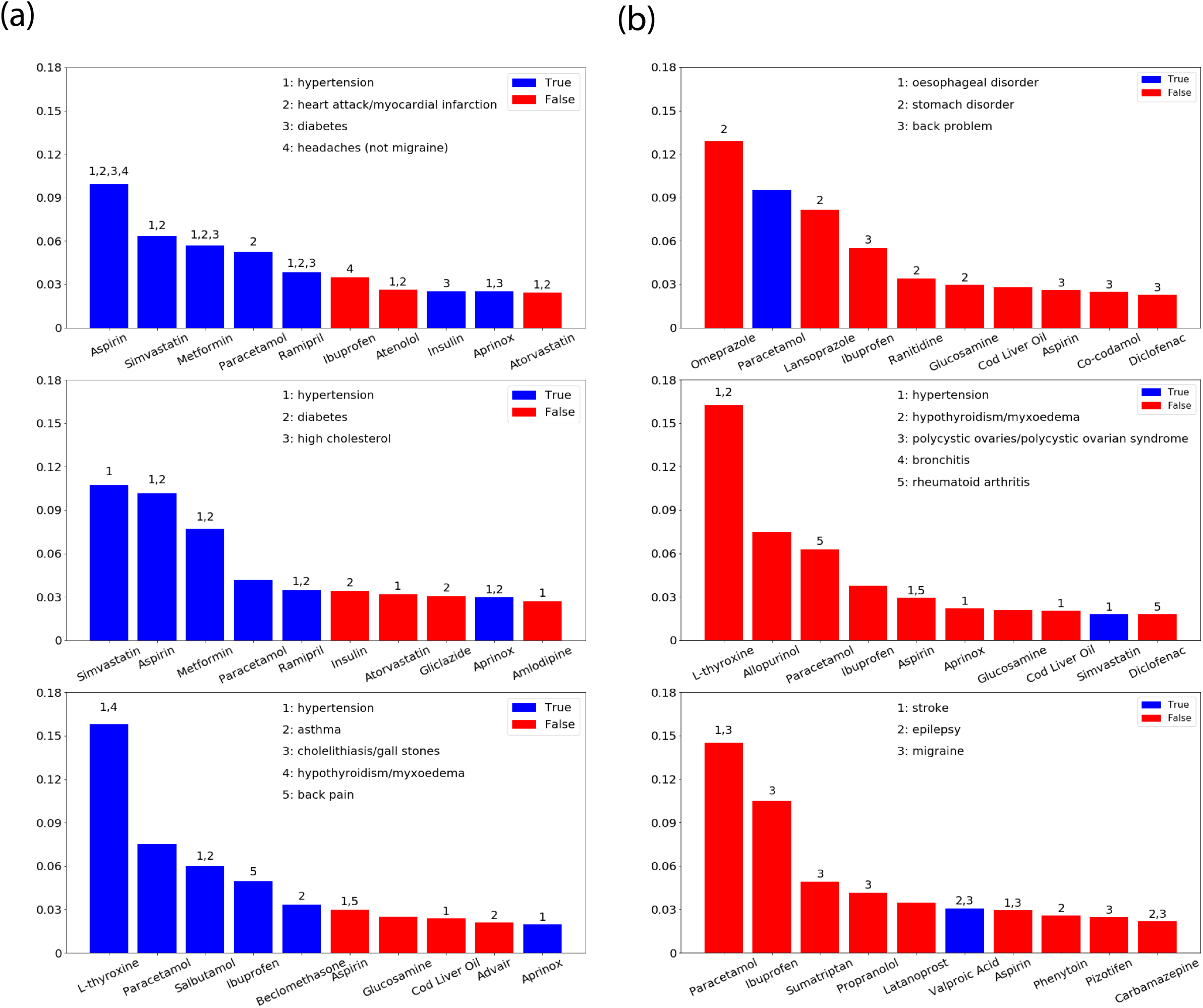
Illustration of GETM for medication imputation through examples of study subjects. (**a**) The three best-matched study subjects displayed as 3 barplot panels on the left. We chose 3 best-matched individuals for whom our imputed medications matched well with the observed medications. (**b**) The three worst-matched individuals displayed as 3 barplot panels on the right. Y-axis represent the predicted probability of the medications on the x-axis. The numbers on the top of each bar represents the patient’s observed conditions that are known to be treated by the corresponding medication under the bar. Inset in each panel lists the conditions names corresponding to the numbers on top of each bar. Blue bars indicate observed medications in the patient, and red bars indicate unobserved medications.

### 2.5 CMK pain prediction

We sought to investigate the predictive ability of reported past conditions and current medications on pain experience in the UKB using GETM. We first focused on CMK pain because it has the highest number of positive cases. Using logistic regression (LR), we evaluated the predictive accuracy of the inferred topic mixture on CMK pain in terms of area under the receiver operating characteristic (AUROC) curve and area under the precision-recall curve (AUPRC). We used GETM with 128 topics for this experiment for the reason of its high performance on the validation data set (**Supplementary Fig. ??**). As a baseline (i.e., the raw model), we trained another LR model that directly used all of the 443 conditions and 802 medications. We also sought to investigate the relative improvements of GETM over standard topic modeling after removing obvious conditions or medications for CMK pain. We experimented 6 different ways of filtering out features based on ORs or expert knowledge (**STAR Methods**).

Predictive performance using the original features (i.e., unfiltered) and the filtered features is summarized in **Fig**. 5. From these results, we observed that (1) The GETM topic mixture (GETM, **Supplementary Table ??**) achieved larger AUROC and larger AUPRC across all feature filtering regimes (i.e., no filter plus the 6 filtering rules; **Supplementary Table ??**); (2) As we removed more pain-related signature conditions and medications, the performance of using raw features dropped more drastically than that using the patient topic mixtures. In other words, the relative improvement of using patient topic mixtures over using raw data increased as we removed more indicative conditions and medications. In particular, GTEM conferred a larger than 40% improvement over the baseline when using the fewest conditions and medications (i.e., m579c322: filtered feature set 7) in contrast to less than 5% improvment over the baseline when using all features (m802c443: filtered feature set 1; **Fig**. 5; **Supplementary Table ??**). These results echo the superior imputation performance of GETM we observed above. This is because GETM uses the pre-trained graph embedding and inferred topic mixture to compensate for the information loss from the feature removals. Nonetheless, the absolute values of AUROC and AUPRC are not high. We discussed these results as limitation of the study in the **Discussion** section.

**Figure 5:**
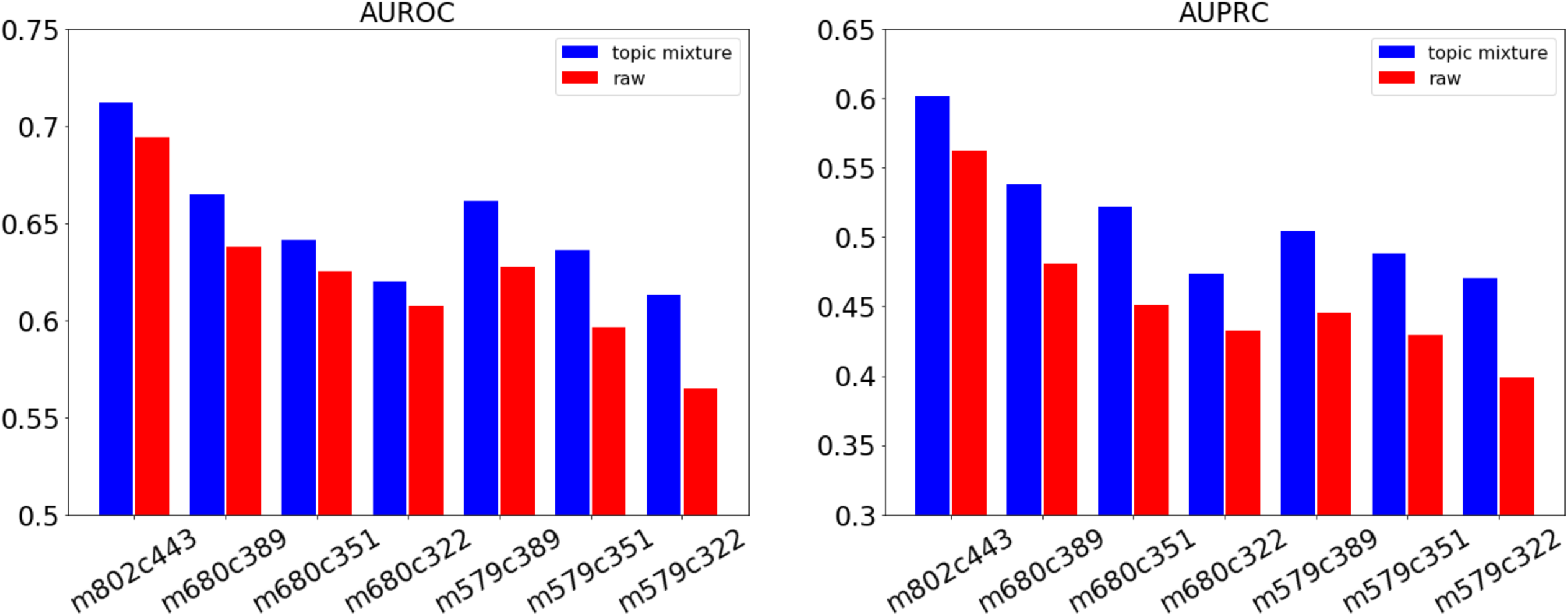
Performance of logistic regression (LR) for CMK pain. We trained LR models using ***θ*** obtained from GETM (**Supplementary Table ??**) with 128 topics as input to predict CMK pain. The baseline LR model directly used the raw condition and medication data as input. We iterated through seven feature filtering schemes with different filtered condition sets and medication sets as indicated by x-axis (details in **Supplementary Table ??** and described in **STAR Methods** Section 8). Barplot displays the AUROC and AUPRC across these experiments.

### 2.6 CMK pain-related conditions and medications

We then investigated the most pain-related conditions and medications based on LR coefficients and calculated overlapping proportions with physician-curated lists (**Fig**. 6). In comparison with the overlapping proportions from ETM and odds ratio calculation, GETM identified a much greater proportion of known pain-related conditions among the top conditions and medications under these topics (36.7% from top 10 conditions, 33.3% from top 30 conditions and 30.0% from top 50 conditions) and medications (60.0% from top 10 medications, 33.3% from top 30 medications and 32.0% from top 50 medications) in the provided lists. This suggests that GETM improves the ability to extract otherwise hidden associations, which could identify pain-related comorbidities. Therefore, GETM does not only confer superior performance but higher model explainability from its inferred topics.

**Figure 6:**
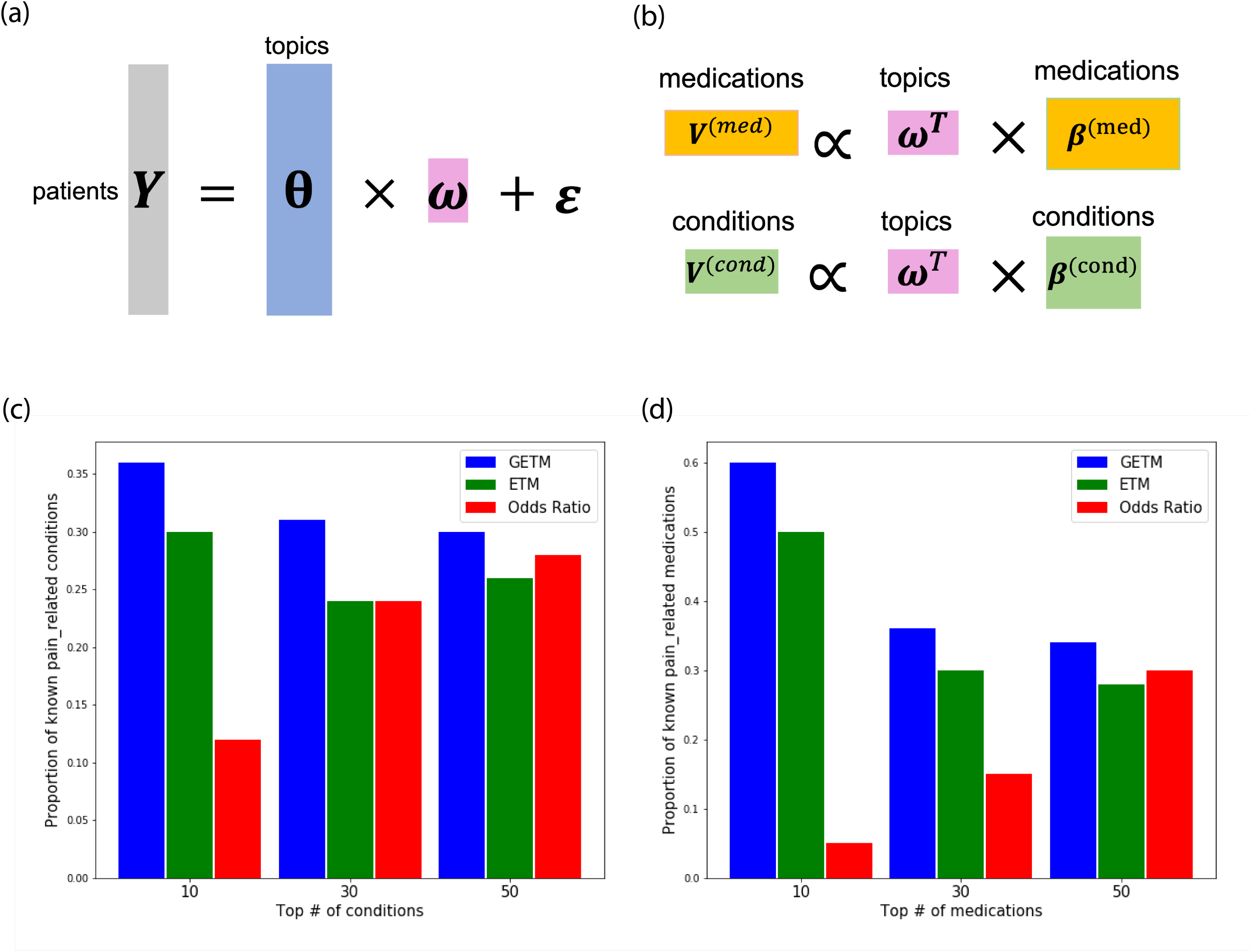
Analysis of CMK-pain-related conditions and medications. (**a**) Logistic regression using patient topic mixture ***θ*** *∈* ℝ^*D×K*^ of D individuals and K topics to predict CMK pain as a binary outcome. **(b)** Importance score computation for medications and conditions. Taking the inner product of the regression coefficients ***ω***^r^ *∈* ℝ^1*×K*^ and ***β***^(*med*)^ *∈* ℝ^*K×M*^ from GETM (GETM, **Supplementary Table ??**), we obtained the importance scores of predicting CMK pain as **v**^(*med*)^ *∈* ℝ^1*×M*^ and **v**^(*cond*)^ *∈* ℝ^1*×C*^ for medications and conditions, respectively. (**c**) Proportions of known CMK-related conditions based on a physician-compiled list. (**d**) Proportions of known CMK-related medications based on a physician-compiled list of analgesic medications. Comparisons relate to two baselines: (1) Using ETM which treated conditions and medications as the same type of feature (i.e., ETM-both-flat, **Supplementary Table ??**). For this baseline, we selected the top medications and conditions from the resulting **v** *∈* ℝ^1*×*(*M* +*C*)^. (2) Odds Ratio (**STAR Methods** Section 8).

We also had a close look at the three most positively associated topics and three most negatively associated topics to CMK pain based on learned ***ω*** (**Fig**. 7a). This analysis showed that topics 56, 34, and 51 are strongly positively associated with CMK pain and topics 73, 68, 89 are strongly negatively associated with CMK pain, respectively. For each topic, we examined their semantic meaning according to the top conditions and top medications (**Fig**. 7b,c). Particularly, topics 56 and 34 contained musculoskeletal system conditions and medications, which are clinically meaningful since they are highly related to CMK pain. In particular, prolapsed disc or slipped disc as a condition is painful, and ibuprofen is an analgesic in the NSAID (non-steroidal anti-inflammatory drug) class.

**Figure 7:**
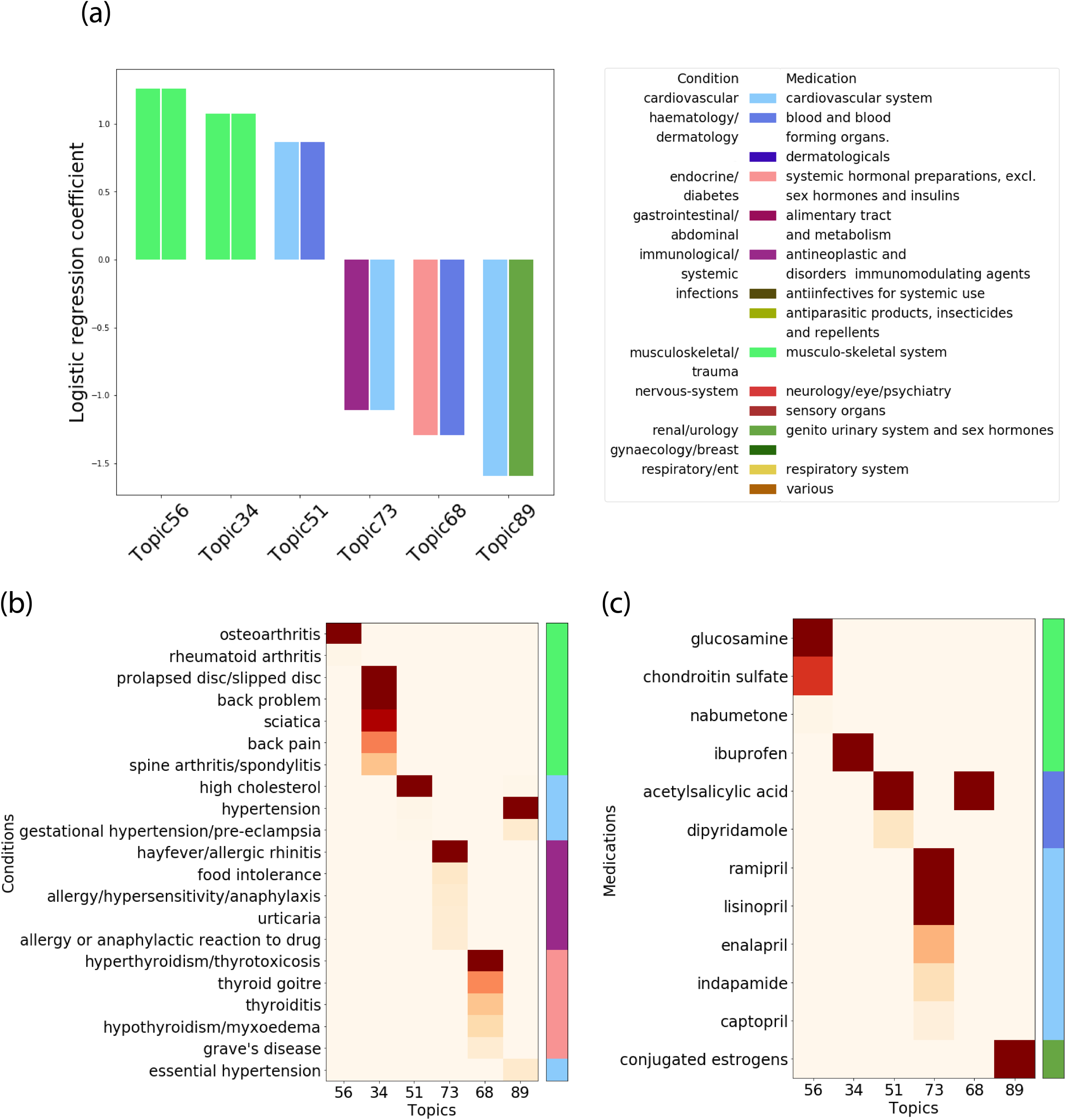
Topic analysis for CMK pain. (**a**) The most predictive CMK pain topics. Based on the logistic regression coefficients of predicting CMK pain ***ω*** (**Fig**. 6a), we chose three topics with the highest coefficients and three topics with the most negative coefficients. Each bar is composed of condition and medication category. (**b**) & (**c**) The top conditions and medications under the 6 most predictive topics. Same as in **Fig**. 2, the color bars on the right indicate the categories of the conditions and medications.

Topic 51 is in the cardiovascular category, and the top medication acetylsalicylic acid (aspirin), also an NSAID, is prescribed more frequently and typically at lower doses for its cardioprotective properties in prevention of stroke and heart attacks; it acts as a “blood thinner”. Dipyridamole inhibits blood clot formation and therefore prevents potential consequences of blood clotting. The top condition under topic 51 is high cholesterol, which is a known and common risk factor for atherosclerosis. Atherosclerosis is a process of deposition of fatty material in the walls of arteries, and this thickening leads to an increase in stroke and heart attack risk. Thus, although the two medications are not directly used as a cure for the condition high cholesterol, by way of atherosclerosis, high cholesterol leads to higher risk for other cardiovascular outcomes and the medications are used prevent those outcomes [**?**].

Contrary to the risk topics, the protective topics identified combinations of medications and conditions that did not yield as straightforward pairings or links. Topic 73 is one of the top 3 negatively predictive topics of CMK pain. Its top medications identified were ramipril, lisinopril and enalapril, which are Angiotensin-converting enzyme (ACE) inhibitors, used to treat high blood pressure and may be used in response to heart failure or heart attack. Incidentally, all conditions under topic 73 may be categoriezed as allergic or atopic with hayfever contributing the most.

This finding suggests that a particular subset of individuals suffering from allergic conditions, whose immunity is therefore skewed, and who are also undergoing cardiovascular treatment, are at lower risk for CMK pain. The implication of immune mechanisms in chronic pain manifestation is well known, where proinflammatory states are associated with chronic pain [**?**]. Also, high blood pressure is the most common indication for ACE inhibitors [**?**]. It is known to have analgesic effects that persist with treatment [**?, ?**]. Thus, hypertension related analgesia would help explain this negative predictive topic.

Another negatively predictive topic of CMK pain, topic 89, is limited to women given the top medication, conjugated estrogens. The top condition, hypertension, as seen above, is known to have analgesic effects. Sex hormones including estrogens modulate pain, and estrogens have been found to have neuroprotective, antinociceptive properties. These could explain the protective association observed here on CMK pain. Indeed, estrogens are known to influence chronic pain conditions [**?**]. These combinations of medications and conditions within topics could lead to new CMK pain etiology hypotheses for further exploration.

### 2.7 Characterizing diverse pain types by topic analysis

We extended our analysis from CMK to predicting several other definitions of pain phenotypes, including acute pain, the acute to chronic pain transition for a subset of UKB individuals, chronic pain at specific body sites (neck/shoulder, hip, back, stomachache/abdominal, knee, headache, face) and chronic pain all over body. We used three different feature filtering regimes to progressively remove obvious pain-type predictors (**STAR Methods**). Logistic regression using the GETM’s topic mixture conferred larger AUROC and larger AUPRC values across all three feature filtering regimes (**Fig**. 8; **Supplementary Table ??**). We then examined the relative contributions of the medication/condition categories for each pain type (**Supplementary Fig. ??**; **STAR Methods**). The results we presented below came from at least one of the 3 feature filtering regimes as highlighted in **Supplementary Fig. ??**. In addition to the defined categories, we used the Anatomical Therapeutic Chemical (ATC) classification system to consider the contribution of the medication sub-class analgesics, which has a direct relationship with pain. By considering the implication of condition and medication categories in predicting different pain phenotypes, we were able to identify expected trends and to consider and interpret unexpected findings.

**Figure 8:**
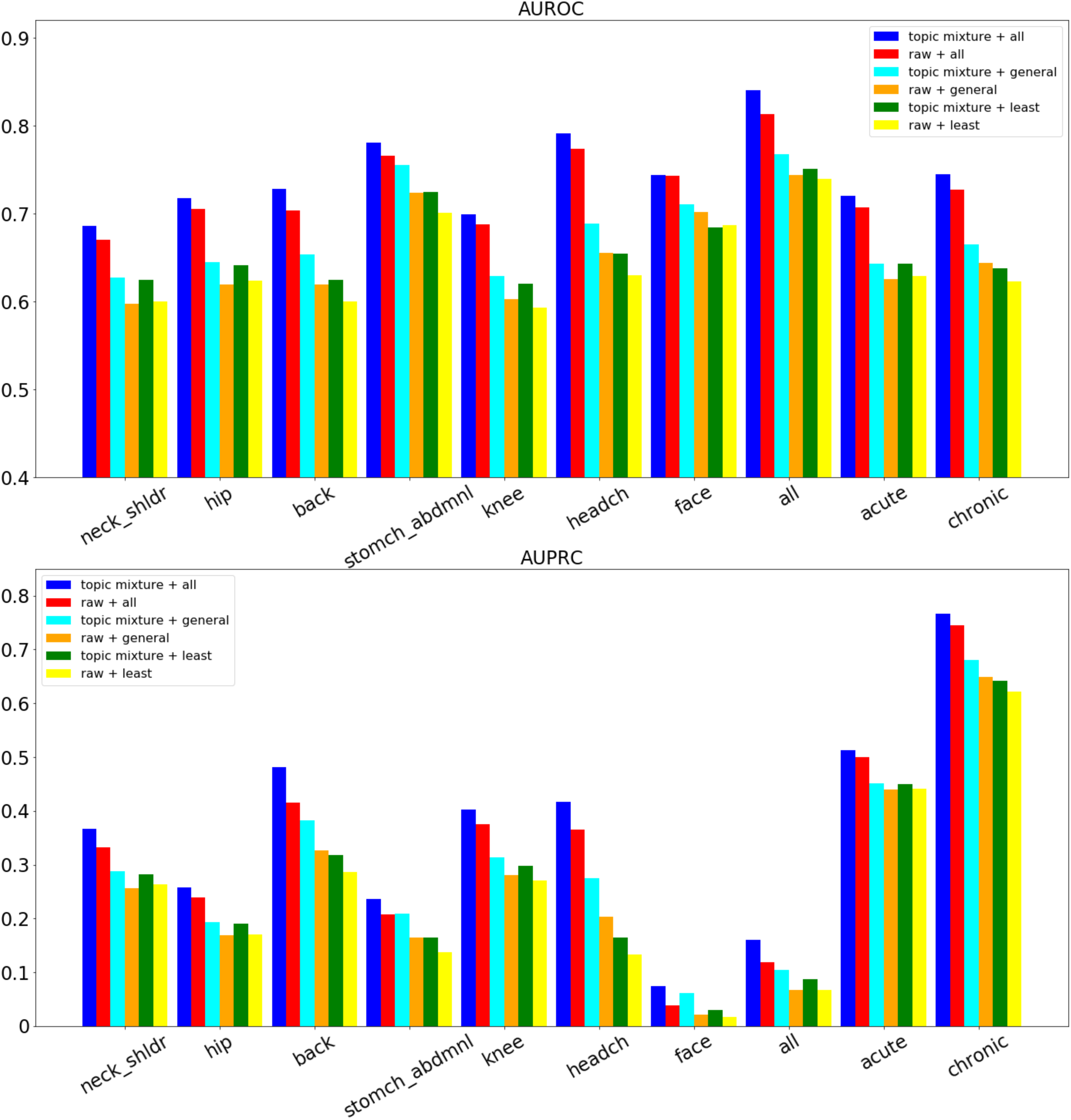
Prediction performance of 10 pain types. Logistic regression was trained to predict 10 pain types as indicated in the x-axis. We experimented with three different filtered feature sets of conditions and medications: (1) <u>all</u> (m802c443): non-filtered conditions and medications, (2) <u>general</u> (m680c351): physician-curated general pain-related conditions and medications were filtered, and (3) <u>least</u>: physician-curated and conditions and medications based on with odds ratios were filtered. Details of the data filtering were described in **Supplementary Table ??**.

Overall, the acute and acute-to-chronic transition displayed similar trends. Here the acute-to-chronic transition is defined such that non-site specific MSK acute pain observed at the first visit turned into chronic pain as diagnosed at the following visit of the UKB individuals. As an example, low positive prediction from analgesics and high negative contribution were seen for acute pain (**Fig**. 9).

**Figure 9:**
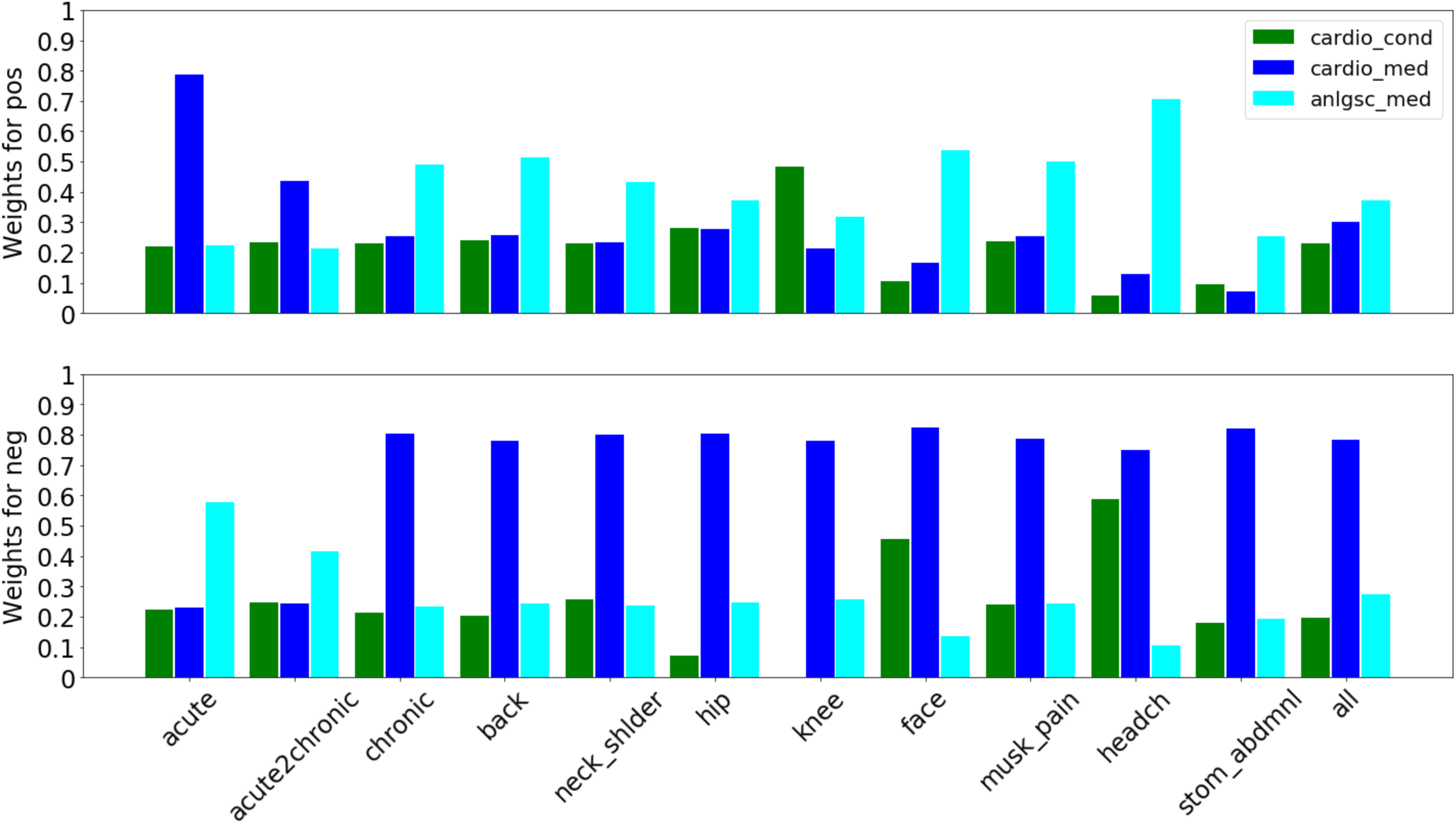
Contributions of cardiovascular-related conditions and medications to 12 pain types. The importance weights of cardiovascular conditions and medications in predicting the 12 pain types were calculated by the sum of their topic probabilities weighted by the linear regression coefficients across all topics (**STAR Methods**). Descriptions of the names of these pain types are in **Supplementary Table ??**. The results for cariovascular category were based on the filtered sets, where the known pain-related conditions and medications based on a physician curated list were removed. The results of analgesic category (anlgsc_med) were based on the non-filtered set, as only full set contains analgesic medications. All of the importance weights from other filtering schemes were displayed in **Supplementary Fig. ??**.

As expected, the category from the literature that displays the highest levels of chronic pain co-morbidities, neurology/eye/psychiatry (conditions) and nervous system (medications) contributed the most to prediction of chronic pain, particularly headache and face pain (**Supplementary Fig. ??**a-I, b-I). Endocrine/diabetes (conditions) exhibited highly prominent contribution for the acute phenotype classification (**Supplementary Fig. ??**a-II). Immunological/systemic disorders showed a strong contribution for protection from acute pain (**Supplementary Fig. ??**c-I). Gastrointestinal/abdominal (conditions) and alimentary tract and metabolism (medications) showed an expected higher positive proportion of contribution towards stomach/abdominal pain predication compared to other chronic pain types (**Supplementary Fig. ??**a-III).

Knee pain compared to face pain and headache exhibited a sharp contrast in predictive compositions (**Supplementary Fig. ??**a, b). Interestingly, obesity-related medication categories such as systemic hormonal preparations excluding sex hormones and insulins exhibited the highest protective contribution for chronic knee pain (**Supplementary Fig. ??**d-I). Chronic knee pain also had the highest positive contribution toward the prediction of cardiovascular condition and no negative contribution (**Supplementary Fig. ??**a-I, c-I). A similar but less striking pattern was observed for hip pain (**Supplementary Fig. ??**a-II, c-II). For headache and face pain, this trend was reversed with the highest protective contribution and lowest positive contribution by cardiovascular conditions (**Supplementary Fig. ??**a-III, c-III).

Intriguingly, the cardiovascular medications category exhibited strong negative prediction of 0.7 across all chronic pain phenotypes but not acute pain (**Fig**. 9). This suggests that topics containing cardiovascular medications, other than analgesics, had a protective effect against chronic pain. Headache, and more specifically migraine, has a known vascular etiological component. Nonetheless, the patterns seen across all chronic pain phenotypes is similar, indicating a more general chronic pain protective effect for cardiovascular medications.

## 3 DISCUSSION

Large biobanks with clinical data such as the UKB are valuable resources enabling greater understanding of factors impacting the manifestation and treatment regimens of complex diseases such as chronic pain. However, the sparsity, heterogeneity and sheer size of these data pose challenges when restricted to a classical statistical toolbox. The typical biomedical researcher is hindered from taking full advantage of these invaluable data resources. As a result, most investigations focus on one or a small subset of related diseases [**?, ?, ?**]. In the present study, we developed GETM to model all self-reported conditions and medications among 457,461 UKB subjects. Our main focus is to understand the phenotypic co-morbidity conditions among the UKB subjects in relation to pain-related phenotypes. By introducing the knowledge graph and simultaneously training different types of features, GETM was able to infer more coherent topics compared to the ablated models without using the KG embedding (**Supplementary Table ??** and **Table** 1). In contrast to the topic modeling without the graph embedding, this allows for better interpretation of the disease topics and any findings related to certain topics with clearer clinical grounds.

**Table 1:**
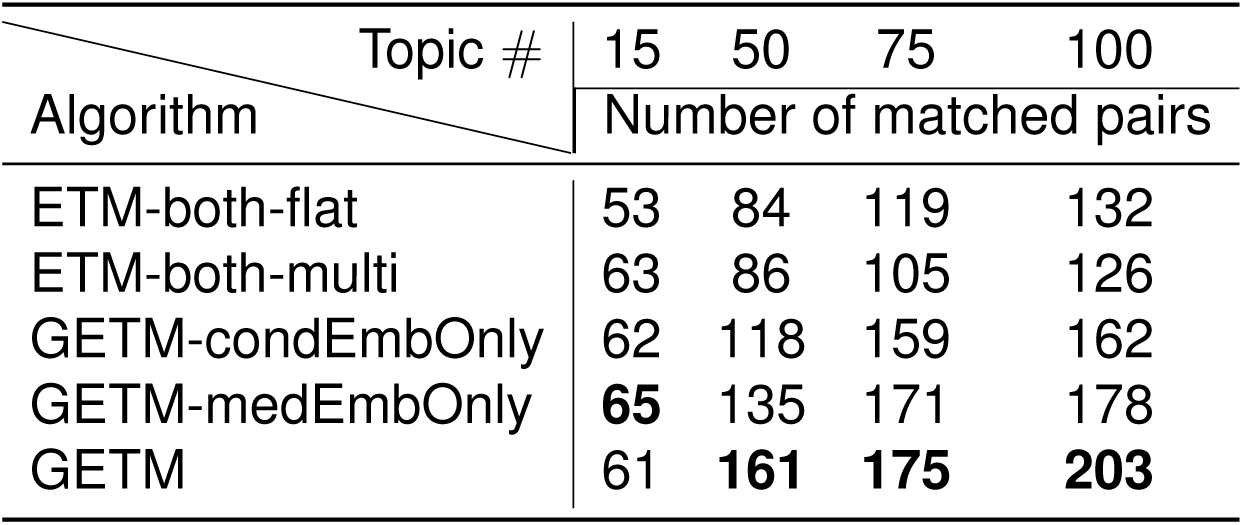
Number of known pairs between conditions and medications. We mapped our medications and conditions identifiers to CTD and DrugBank databases to obtain known connections between them. For each topic inferred, we generated 9 condition-medication pairs from the top 3 conditions and the top 3 medications. Among these pairs, we calculated the number of known pairs (**Supplementary Table ??**). Higher numbers imply more clinically meaningful topics inferred by the algorithm.

As demonstrated using UKB data, GETM achieved superior performance in imputing 50% of the observed conditions (**Table** 2) and 50% of the observed medications over all test individuals (**Table** 3) as well as in imputing the entire medication records based only on the conditions records of test individuals (**Table** 4). Many top unobserved medications predicted by GETM also exhibit known connections with the observed conditions of the subject (**Fig**. 4). Importantly, GETM offers excellent model interpretability and can be used to discover meaningful disease comorbidities and disease-medication combinations via topic analysis (**Fig**. 2).

**Table 2:** Reconstructing 50% masked conditions. We split the UKB data into 80% training and 20% test. For the test data, we randomly masked 50% of the values such that each test patient would have 50% of their conditions and medications observed. We then reconstructed the matrix with learned ***θ, α*** and ***ρ***. The reconstruction error (i.e. negative log-likelihood) was calculated for the held-out data. Same condition data was used for all algorithms. For ETM-both-multi, GETM-condEmbOnly, GETM-medEmbOnly and GETM, the same models and data were used as for the medication reconstruction **Supplementary Table ??**. Description of the algorithm names are in **Supplementary Table ??**. As another baseline method, we evaluated the performance of filling in masked conditions based on their overall proportion over all of the UKB population.

| Algorithm \ Topic # | 15 | 50 | 75 | 100 |
| --- | --- | --- | --- | --- |
|  | Reconstruction Error |  |  |  |
| ETM-condOnly | 5.89 | 5.56 | 5.39 | 5.80 |
| GETM-condOnly | 5.76 | 5.33 | 5.05 | 4.93 |
| ETM-both_multi | 5.08 | 5.09 | 4.80 | 4.96 |
| GETM-condEmbOnly | 5.12 | 4.84 | 4.55 | 4.40 |
| GETM-medEmbOnly | 5.06 | 4.86 | 4.75 | 4.58 |
| GETM | **4.69** | **4.84** | **4.45** | **4.34** |
| Proportion | 8.69 |  |  |  |

**Table 3:** Medication imputation. The medication data was masked for test individuals. In contrast to the medication reconstruction experiments (**Supplementary Table ??**), we masked the entire medications for the test patients. We imputed their medication data using the inferred ***θ*** from their condition data only and the learned embedding ***α***^(*med*)^ and ***ρ***^(*med*)^. The reconstruction error (i.e. negative log-likelihood) was calculated. The upper bounds were obtained using reconstruction errors calculated from unmasked test data using GETM (GETM, **Supplementary Table ??**). Description of the algorithm names are in **Supplementary Table ??**.

| Algorithm \ Topic # | 15 | 50 | 75 | 100 |
| --- | --- | --- | --- | --- |
|  | Medication Recovery Error |  |  |  |
| Upper bound | 12.25 | 11.24 | 11.50 | 11.50 |
| ETM-both-flat | 18.51 | 19.85 | 19.81 | 20.06 |
| GETM-both-multi | 14.99 | 14.88 | 14.94 | 15.05 |
| GETM-condEmbOnly | 15.24 | 14.95 | 14.95 | 14.96 |
| GETM-medEmbOnly | 15.18 | 14.88 | 14.90 | 14.88 |
| GETM | **14.82** | **14.65** | **14.61** | **14.65** |

**Table 4:** Medication imputation accuracy. The imputation procedures were the same as in **Table** 3. We calculated the precision and recall at the top 5 imputed medications for each individual. The upper bounds were calculated from unmasked test data using GETM (GETM, **Supplementary Table ??**).

| Algorithm \ Topic # | 15 | 50 | 75 | 100 | 15 | 50 | 75 | 100 |
| --- | --- | --- | --- | --- | --- | --- | --- | --- |
|  | recall@5 |  |  |  | precision@5 |  |  |  |
| Upper bound | 0.6672 | 0.7943 | 0.8027 | 0.7862 | 0.3342 | 0.4084 | 0.4226 | 0.4049 |
| ETM-both-flat | 0.4397 | 0.4486 | 0.4667 | 0.4614 | 0.1901 | 0.2109 | 0.2111 | 0.2162 |
| GETM | 0.5543 | 0.5639 | 0.5568 | 0.5388 | 0.2479 | 0.2516 | 0.2504 | 0.2417 |
| GETM-condEmbOnly | 0.5479 | 0.5664 | 0.5670 | 0.5647 | 0.2373 | 0.2524 | 0.2493 | 0.2486 |
| GETM-medEmbOnly | 0.5519 | 0.5722 | 0.5668 | 0.5716 | 0.2440 | 0.2533 | 0.2521 | 0.2532 |
| GETM | **0.5692** | **0.5787** | **0.5732** | **0.5753** | **0.2504** | **0.2612** | **0.2606** | **0.2578** |

In a focused analysis to predict chronic musculoskeletal (CMK) pain, the same logistic regression (LR) classifer using GETM-inferred topic mixtures conferred robustly and consistently higher AUROC and AUPRC compared to using the raw features (**Fig**. 5). The top conditions and medications under the most predictive topics are enriched for elements from the physician-curated condition list and medication list, respectively (**Fig**. 6). Also, the most predictive CMK pain topics contained top conditions and medications that were strongly associated with CMK pain (**Fig**. 7), implying their potential usage as phenotyping markers. Additionally, the GETM topic mixtures are robust to the removal of clinically obvious conditions and medications (**Fig**. 5). This is reflected in the larger relative improvements over the LR classifer that operates on the raw (filtered) features. This implies its utility in predicting predisposition for CMK pain for those individuals with no reported pain symptoms. We observed consistent quantitative performance when extending to other pain types (**Fig**. 8).

Several existing methods utilize topic models to find meaningful latent topics from electronic health record (EHR) data using structured administrative data such as the ICD codes [**?, ?**]. In contrast, we demonstrated the utility of GETM on the less structured and more sparse self-reported questionnaire information from the UKB including 443 conditions and 802 medications. We expect that GETM would work equally well if not better on more structured EHR data and leave that to future exploration. Indeed, although we used UKB data and focused on pain phenotypes as a case study, our approach is a generalizable and highly efficient method that can be used to characterize other phenotypes in the UKB or from other similarly scaled biobanks.

We now discuss the epidemiological implications of our results on the pain analysis in the context of existing studies. Our findings identified intriguing links between cardiovascular medications and their protective effect for chronic (but not acute) pain that stood out as particularly predictive compared with other medication or condition categories (**Fig**. 9). Also, cardiovascular conditions were negatively predictive for headache and face pain particularly. Recent meta-analyses have attempted to quantify the relationship between cardiovascular conditions and chronic pain [**?, ?**], but their relevance to our findings is low. None of these, nor the studies that were included in the meta-analyses explicitly quantified the effects of taking medications. Also, the direction of causality focused on chronic pain as a risk factor for cardiovascular outcomes, including mortality, and did not consider reverse causality; authors cited diversity in outcomes and in chronic pain taxonomy making meta-analysis results generally inconclusive [**?**]. The more recent meta-analysis was more limited in scope, estimating that people with CMK pain were 1.91 times more likely to report having a cardiovascular disease compared with those without CMK pain with statistical significance [**?**].

The dominance of the predictive effect of cardiovascular medications across chronic pain phenotypes leads to several avenues for further exploration. The specifics of what medications and what conditions are found in individually highly predictive topics would need to be explored in further detail. Also, further analyses are needed to identify whether more specific sub-categories of cardiovascular medications drove the strong cardiovascular medications predictive component. The extent of similarity across specific medications within and across topics, such as common mechanisms of action, would be of particular interest. Several medications united by a common mechanism would have a higher potential for generalizability beyond the limited population subsets represented by individual items or topics. This was illustrated by our focused analysis on CMK pain, where the predictive medications identified in the top protective topic (**Fig**. 7b,c, Topic 73) were all ACE inhibitors. Given that the most common indication for this medication class is hypertension, perhaps one of the protective cardiovascular medication effects at play was hypertension-associated hypoalgesia, especially since the hypoalgesia effect is maintained even if the hypertension is treated [**?, ?**]. Baroreceptor sensitivity is the most studied specific mechanism [**?, ?**]. Hypertension-associated hypoalgesia manifests through a range of phenotypes from pain sensitivity to pain chronification, with higher blood pressure associated with lower pain sensitivity or lesser chronification. This has been demonstrated in both animal experimental and human observational studies [**?, ?**]. Nonetheless, hypertenstionassociated hypoalgesia is not well recognized.

Perhaps the present report will lead to findings supporting greater clinical and public health importance for hypertension-associated hypoalgesia in preventing pain chronification than previously thought. Beyond ACE inhibitors, cardiovascular medications classified as beta blockers are used as prophylaxis for migraine headaches [**?**]. They have been shown to have analgesic effects for other types of chronic pain [**?, ?**]. There may also be further sub-categories of cardiovascular medications of interest.

### 3.1 Limitations of the study

In terms of the data used in this study, the UKB as a cross-sectional study creates challenges in interpreting the role of temporality and identifying putative causal effects. The pain questionnaire was administered at baseline, and medications reported were taken regularly, also at baseline. Conditions were considered to occur at any time in the subjects’ past, through a record of age-of-onset for each condition. Thus, the impact of medications on reported conditions do not allow for direct investigations of medication efficacy. In classifying chronic pain, the pain questionnaire referred to a time including at least the prior three months in the past from baseline, but this could extend to any length in a given subject’s history. By considering the subset of individuals without baseline chronic pain, but who developed chronic pain by the time of a subsequent visit, it would be possible to approximately quantify the total length of time of pain chronicity. We may also consider chronification as temporally subsequent to the appearance of conditions and taking of medications from prior to baseline. This is only true for a subset of UKB subjects, however. Thus, the UKB study design has limited potential to evaluate chronic pain causality. Rather, our analyses produced hypotheses that might best be tested in the context of other cohorts that are truly longitudinal in design.

In terms of the biological findings in this paper, there are variable time frames between development of cardiovascular conditions, taking of cardiovascular medications and development of chronic pain in the present study. Therefore, other longitudinal studies that control these sources of variability would be needed to better understand the protective effect suggested by the present analysis.

Although GETM provides relatively the highest chronic musculoskeletal (CMK) pain prediction accuracy (**Fig**. 5), the absolute AUROC and AUPRC are not high. This underscores the limitation of both the UKB data and the GETM model. First of all, we only used self-reported conditions and medications in this task, which are noisy, sparse, and sometimes inaccurate. To make more accurate predictions on CMK pain, other sources of health status data such as healthcare provider-facilitated clinical notes, laboratory tests, genomic measurements such as gene expression profiles of each patient are needed.

Another important missing piece for predicting chronic conditions is the longitudinal information of each patient. As mentioned above, we only have the baseline data at the initial visit for most of the UKB subjects. As longitudinal data become increasingly available, we will extend our model to a dynamic topic model that accounts for the evolution of subjects’ health status over time. In particular, we will allow for integration of longitudinal healthcare information such as age-of-onset across conditions and multiple follow-up timepoints. We will also improve embedding learning by considering a more expressive graph embedding approach than node2vec. For instance, we will consider a graph convolutional neural network (GCN) [**?**] to produce the embedding used by the ETM model.

Furthermore, we will explore a multi-relational graph approach [**?**] to model the relationships within and among conditions and medications. Since both the GCN and the ETM models share the same objective function (i.e., the evidence lower bound), we will be able to perform end-to-end training instead of the pipeline approach presented here. To produce competitive performance with this approach, however, careful model fine-tuning of both the GCN and ETM will be needed at the expense of computational resources.

## 4 Acknowledgements

Y.L. is supported by Natural Sciences and Engineering Research Council (NSERC) Discovery Grant (RGPIN-2019-0621), Fonds de recherche Nature et technologies (FRQNT) New Career (NC-268592), and Canada First Research Excellence Fund Healthy Brains for Healthy Life (HBHL) initiative New Investigator start-up award (G249591). AVG and YW were partly funded by the Canadian Excellence Research Chairs (CERC09). LD was supported by the Canadian Excellence Research Chairs fund (CERC09), a Pfizer Canada Professorship in Pain Research, and CIHR (SCA-145102) for Health Research’s Strategy for Patient-Oriented Research (SPOR) in Chronic Pain. The current study was conducted under UK Biobank application 20802.

## 5 Author contributions

These authors jointly supervised this work: Audrey V. Grant and Yue Li. Y.L. and A.V.G. conceived the study. Y.L. and Y.W. developed the methodology. Y.W. created the computational software and ran the analyses. Y.W., Y.L., and A.V.G. analyzed the results and wrote the initial manuscript. L.D. and R.B. participated in initial discussions and subsequently in critical writing related to pain phenotypes and medications. All of the authors wrote the final version.

## 10 STAR★METHODS

### 10.1 RESOURCE AVAILABILITY

#### Lead contact

Requests for data and requests for additional information should be directed to the lead contact, Yue Li.

#### Material availability

This study did not generate new unique reagents.

#### Data and code availability

- The UK Biobank data access has been approved by McGill IRB under the project title “A replication study of pain interactions with comorbidities”. The approval number is A03-M20-21B.
- All code associated with this paper can be freely accessed and downloaded via https://github.com/li-lab-mcgill/getm
- Any additional information required to reanalyze the data reported in this paper is available from the lead contact upon request.

### 10.2 METHOD DETAILS

#### UKB data processing

For conditions data, we used UKB datafield 20002, which records self-reported non-cancer diseases for each subject. This was collected by questionnaire during participant interviews. The participants were asked whether or not they had been diagnosed with certain conditions (heart attack, angina, stroke, high blood pressure, blood clot in leg, blood clot in lung, emphysema/chronic bronchitis, asthma or diabetes), were asked to add any other conditions and to provide a date of diagnosis for each when possible [**?**]. Medication usage data was similarly collected during participant interviews prompted by a question on regular prescription medication use, resulting in datafield 20003 which contains treatment/medication codes [**?**].

We kept 457,461 individuals of European descent to reduce potential confounding by ethic group. In total, 802 active ingredients were kept as medications and 443 conditions were extracted. Here we encoded the medications and conditions as binary variables. Only baseline visit data was included.

#### UKB pain-related phenotype labels extractions

We sought to associate pain phenotypes with comorbid conditions and medications. To this end, we used the pain-related phenotypes as the label data and patient-dependent topic mixture derived from the 443 conditions and 802 medications as the input features in a post-hoc supervised topic analysis as detailed in Section 2.5.

The pain phenotypes were collected through questionnaire. We drew on datafield 6159; participants were asked “In the last month have you experienced any of the following that interfered with your usual activities? (You can select more than one answer).” If they answer “yes” to pain at any site, for example, back pain, they were further asked “Have you had back pains for more than 3 months?” (datafield 3571), and so on. In total, we collected 9 pain-related labels with data field identifiers listed as follows:

1. pain type(s) experienced in last month: 6159
2. headaches for 3+ months: 3799
3. facial pains for 3+ months: 4067
4. neck/shoulder pain for 3+ months: 3404
5. back pain for 3+ months: 3571
6. stomach/abdominal pain for 3+ months: 3741
7. hip pains for 3+ months: 3414
8. knee pains for 3+ months: 3773
9. general pain for 3+ months: 2956

We can also query the UKB website to obtain relevant information for each field by replacing *x* in https://biobank.ndph.ox.ac.uk/ukb/field.cgi?id=x with the above data field identifiers.

#### Graph-embedded topic model details

##### Graph-embedded topic model (GETM) generative process

We formulated the problem of modelling discrete patient healthcare data into a topic modeling problem. In particular, we treat each of the *D* patient health records as a document and each observed feature in the record as a word sampled from a defined vocabulary. In our case, we have two vocabularies covering *M* = 802 medications and *C* = 443 conditions, respectively. With this analogy, each patient record is represented as a mixture of latent topics. In the original latent Dirichlet allocation (LDA) model [**?**], a topic distribution over a vocabulary is defined as an independent Dirichlet prior ***β***_*k*_ *∼ Dirichlet*(***τ***_***β***_), where 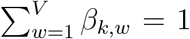 over V words. Inspired by a more recent work called embedded topic model (ETM) [**?**], we decomposed the *K* topic distributions over medications ***β***^(*med*)^ into a medication-defined topic embedding 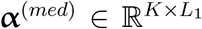, and a medication embedding 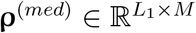, where *L*_1_ denotes the medication embedding dimension and *M* denotes the number of unique medications. Similarly, the topic distribution over conditions ***β***^(*cond*)^ is proportional to the inner product of the condition-defined topic embedding 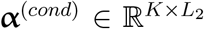, and condition embedding 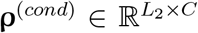, where *L*_2_ denotes the condition embedding dimension and *C* denotes the number of unique conditions.

For the *d*^*th*^ patient record (*d ∈ {*1, …, *D}*), the generative process starts by drawing the topic mixture ***θ***_*d*_ from a logistic normal distribution ***θ***_*d*_ *∼ ℒ𝒩* (0, **I**):

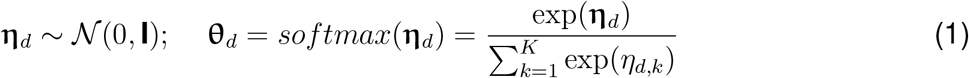

We then draw the *i*^*th*^ medication token or the *j*^*th*^ condition token from two respective categorical distributions:

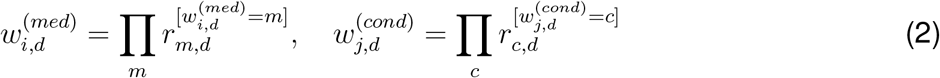

where

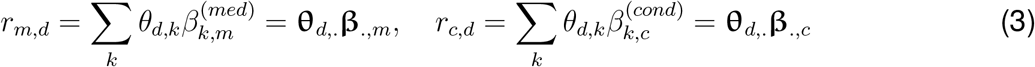

Here we use notation ***θ***_*d*,._ to denote a 1 *× K* row vector for the *d*^*th*^ patient record and ***β***_.,*m*_ (and ***β***_.,*c*_) to denote a *K ×* 1 vector for the *m*^*th*^ medication (and the *c*^*th*^ condition).

The *k*^*th*^ topic distribution for the *m*^*th*^ medication (or the *c*^*th*^ condition) is defined as the product of the corresponding topic embedding and medication (or condition) embedding followed by softmax normalization over the corresponding vocabularies:

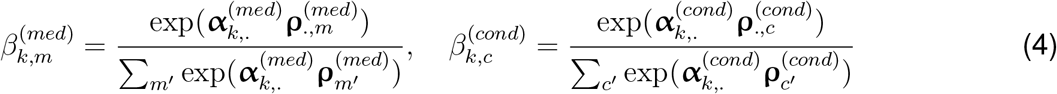

Here we treat both the topic embeddings ***α***^(*med*)^ and ***α***^(*cond*)^ and condition and medication embeddings ***ρ***^(*med*)^ and ***ρ***^(*cond*)^ as point estimates without imposing any prior distribution.

For the ease of mathematical expression below, we denote 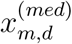 (and 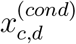 as the frequency of the *m*^*th*^ medication (and the *c*^*th*^ condition) for the *d*^*th*^ patient record. Formally, modeling the likelihood of the count frequency simply requires reformulating the above multinomial likelihood:

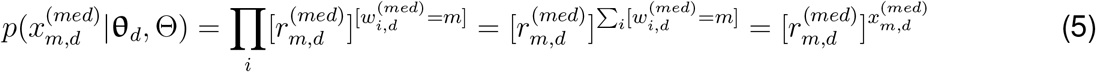

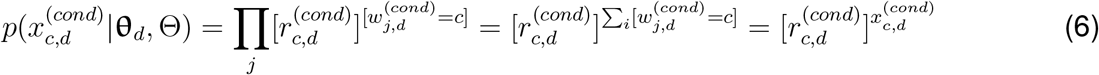

where ***θ***_*d*_ is the patient topic mixture of patient *d* and Θ = *{****α***^(*med*)^, ***ρ***^(*med*)^, ***α***^(*cond*)^, ***ρ***^(*cond*)^}.

The vector 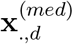 and 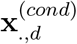 denote the frequency of all medications and all conditions for the *d*^*th*^ patient record, respectively. The entire data can then be represented as a *M × D* matrix **X**^(*cond*)^ and a *C × D* matrix **X**^(*cond*)^ and modelled by Eq (5) and Eq (6):

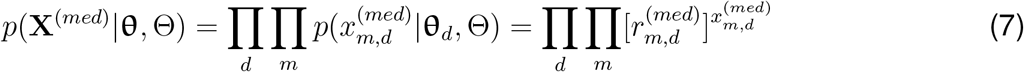

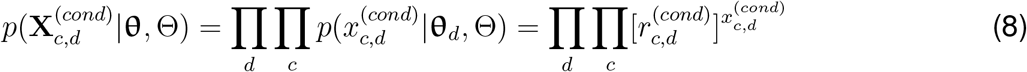

Inference of ***θ***_*d*_’s and learning of Θ are described next.

#### Model inference and estimation

To train GETM, we want to maximize the marginal likelihood of the individuals with respect to Θ = *{****α***^(*med*)^, ***ρ***^(*med*)^, ***α***^(*cond*)^, ***ρ***^(*cond*)^}:

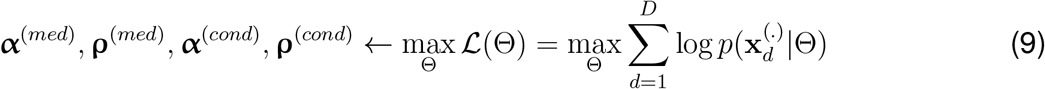

where 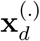 is word frequency vector (i.e., the bag of words of medications and conditions for individual *d*). This marginal likelihood is intractable to compute since it involves integrating out the logistic normal topic mixture variable ***θ***_*d*_, *∀d ∈ {*1, …, *D}*:

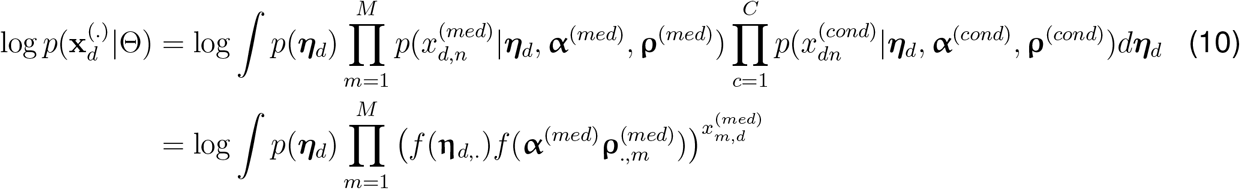

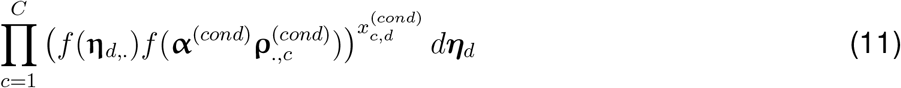

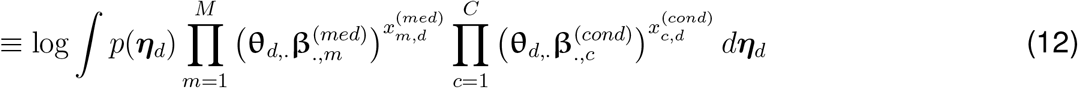

where *f* (.) denotes the softmax function and 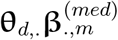 is the inner matrix product of the 1 *× K* vector ***θ*** and the *K ×* 1 vector 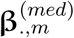. Same for conditions.

Therefore, we took an *Amortized Variational Inference* (AVI) approach to approximate the above intractable integral. This is quite similar to the one described by Kingman & Welling 2014 [**?**] and by Dieng et al 2019 [**?**]. For the sake of completeness, we describe it below.

To approximate true posterior *p*(***η***_*d*_|***x***_***d***_), we define the proposed distribution as a Gaussian with mean and variance produced by a feedforward neural network with parameters **W**_*θ*_:

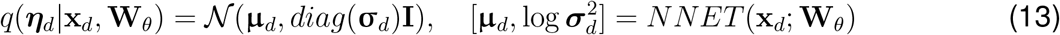

The evidence lower bound (ELBO) of the above log-likelihood is approximated as follows:

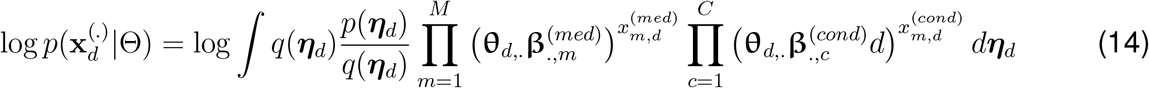

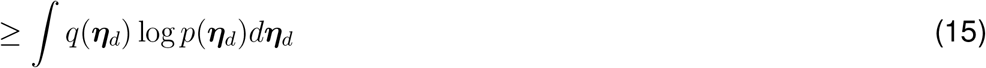

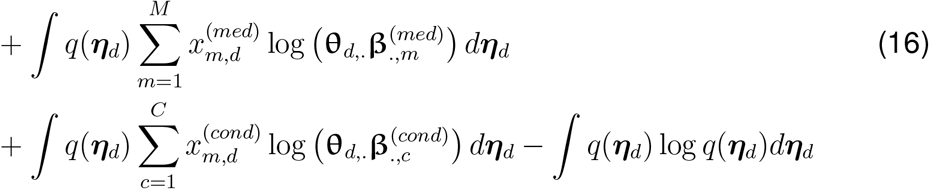

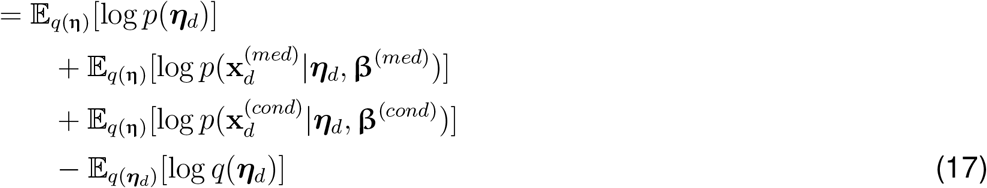

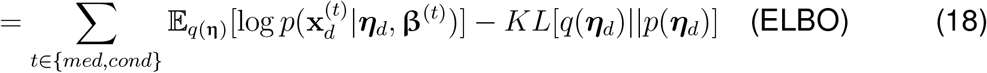

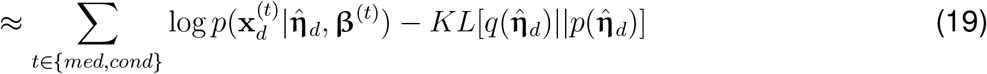

Where

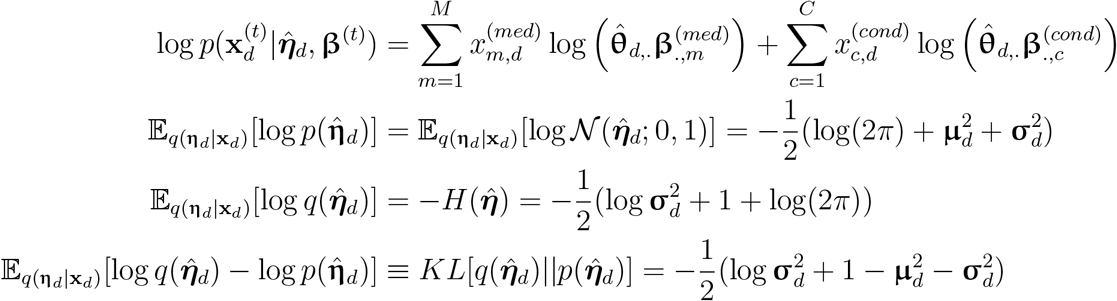

Here Eq (19) uses the sampled 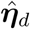 from the feedforward network encoder as a surrogate to approximate the intractable variational expectation w.r.t. 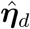. We re-parameterized the Gaussian distribution in Eq (13) to enable stochastic gradient calculation of the ELBO w.r.t. **W**_*θ*_ and AVI training of the encoder network *NNET* :

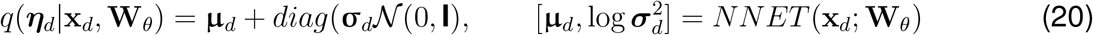

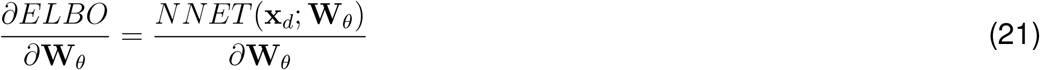

#### Model specifications

The encoders for GETM, partial GETMs and ETMs are 2-layer neural networks with hidden sizes of 128, ReLU activations [**?**] and 1D batch normalization [**?**]. The medication embedding dimension and condition embedding dimension were both 128 and they were fixed during training of the models with KG-informed pre-trained embeddings. The models were optimized with Adam optimizer at 0.01 learning rate. We trained each model with batch size of 100. For inferring individual-topic mixtures used by prediction tasks, the topic number was set to be 128.

#### Knowledge graph construction for learning the embedding of UKB conditions and medications

In the original ETM [**?**], the word embedding ***ρ*** can be either learned from the documents or pre-trained on a separate large data corpus such as Wikipedia using the word2vec approach. The latter approach allows leveraging the contextual data to improve the topic modeling. As one of our main contributions, we sought to develop a simple framework that exploits ETM’s ability to incorporate a pre-trained medication embedding ***ρ***^(*med*)^ and condition embedding ***ρ***^(*cond*)^ from their taxonomic knowledge graphs (KG). Leveraging the structural KG information and the internal relational information among medications and conditions in a principled way is beneficial to modeling the UKB phenotype data and other EHR data in general, which are often sparse, noisy, and bias.

Among the graph representational learning methods, we chose node2vec [**?**] as it is an unsupervised approach that can directly learn from the KG without linking to a prediction task as in other methods. Specifically, we applied node2vec to separately learn the node embedding of the 443 conditions or 802 medications from their hierarchical trees (**Fig**. 1).

For the KG of the 443 conditions, we constructed a tree graph using coding taxonomy designed by the UKB team (i.e., datafield 20002). The tree describes the topology of the conditions with 473 nodes and 4 hierarchical levels.

For the KG of the 802 medications, we constructed a KG based on the Anatomical Therapeutic Chemical (ATC) classification system [**?**]. The entire tree is composed of 5 levels. We first kept the top 4 levels of ATC, of which the first level contains main anatomical or pharmacological groups; the second level includes pharmacological or therapeutic subgroups; and in the third and fourth levels are chemical, pharmacological or therapeutic subgroups. To harmonize the UKB medications with the ATC graph, we mapped the names of active ingredients from UKB datafield 20003 to the fifth level codes of ATC, which are chemical substances. Some UKB medication codes were mapped to multiple ATC fifth level codes as they could belong to different subgroups depending on their usages. In that case, we replaced all of the mapped ATC fifth level codes with the same corresponding UKB medication code of the active ingredient in the second step. The final medication graph contains 2561 nodes in total.

Both the condition graph and the medication graph were treated as undirected graphs as input to the node2vec model (**Fig**. 1).

#### Summary of the GETM learning algorithm

GETM learning algorithm is summarized in Algorithm 1.

##### Algorithm 1: Topic modelling with GETM

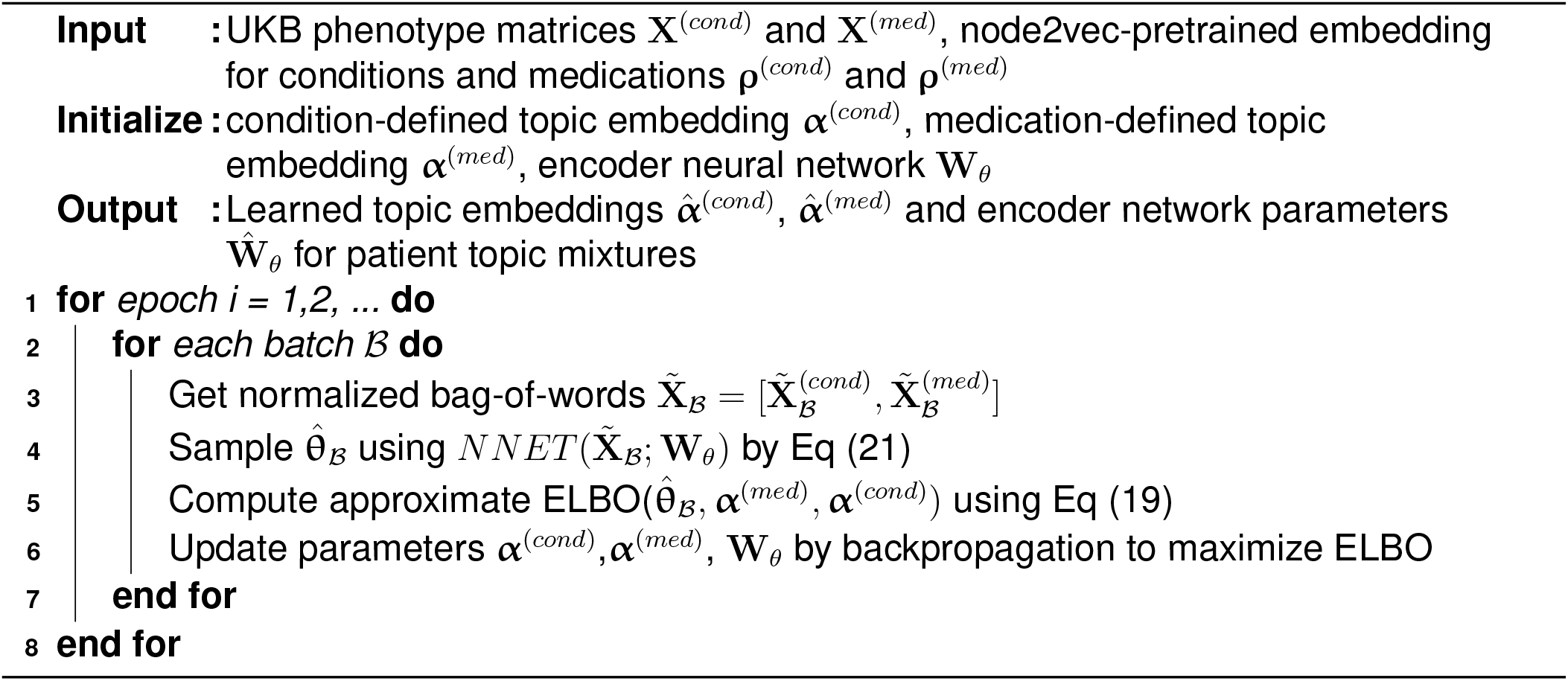

#### Ablation experiments

To gain a good understanding of how each component of our method contribute to the overall improvements, we performed ablation experiments. In particular, we compared GETM with 8 ablated baseline models (**Supplementary Table ??**):

#### Topic quality evaluation

For medication, the topic coherence was calculated as:

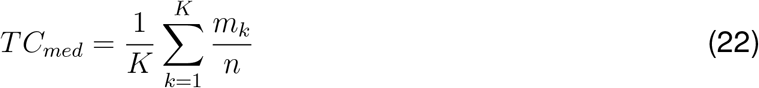

where *n* is the number of top medications, *m*_*k*_ ≤ *n* is the maximum number of medications that are from the same category across all categories, and *K* is the number of topics. We set *n* = 5 because of the softmax-normalization of each topic, where only the top 5 conditions or medications under each topic has non-negligible probabilities.

To avoid overestimating the topic quality, the categories we used to evaluate the topic coherence were not processed from the ATC graph since it was involved in the pre-trained model. Instead, we employed 59 categories which are physician-curated and pain-focused (**Supplementary Table ??**).

For conditions, we did not have any external gold-standard reference to compare against. Therefore, we calculated the topic coherence for conditions based on the co-occurence of the top conditions under the same topic observed from the same patient record. Specifically, for any top two conditions under the same topic, we calculated the average pointwise mutual information:

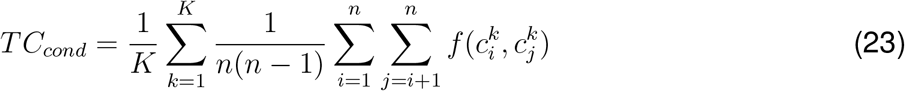

where 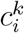 is the *i*^*th*^ top most likely condition in topic k, and *f* (*·, ·*) is normalized pointwise mutual information:

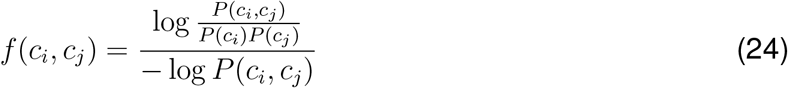

where *P* (*c*_*i*_, *c*_*j*_) is the probability of condition i and condition j co-occurring in one individual and *P* (*c*_*i*_) is the marginal probability of condition i. The probabilities were approximated by empirical counts.

Topic diversity is the percentage of total unique features of the top 5 features of all topics. We first chose top 5 features from K topics. Among 5 *× K* features there are *U* unique words. The topic diversity is calculated as 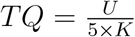. Therefore, the higher topic diversity the more diverse the topics are.

#### Study of medication and condition relations

In GETM, since the encoder takes both medication and condition information as input (**Fig**. 1), the top medications and conditions from the same topic (i.e. the same index) are related. To quantitatively measure the ability of the model to capture known condition-medication relations, we first obtained all of the combinations from top 3 conditions and top 3 medications for each topic. We then counted the number of condition-medication pairs which are known to be related. The reference of known pairs was extracted from CTD http://ctdbase.org/ and DrugBank https://go.drugbank.com/. We eventually mapped 222 conditions and 529 medications from UKB to these two databases. As a result, we had 2444 positive pairs of which the medication has treatment effects on the condition and 3231 negative pairs of which the condition belongs to the adverse effects of the medication. The number of known pairs discovered by our model and baselines were compared using different number of topic numbers (**Table** 1).

#### Data imputation

Using the generative model, we can (1) impute missing entries including conditions or medications and (2) impute medications based on only clinical conditions. Accordingly, two sets of experiments were performed to evaluate how well our model could complete these two tasks. We split the UKB data into 80% training and 20% test. To simulate missing entries, we randomly masked 50% of medications and conditions for test individuals. Then we calculated reconstruction error as the negative log-likelihood of the held-out data:

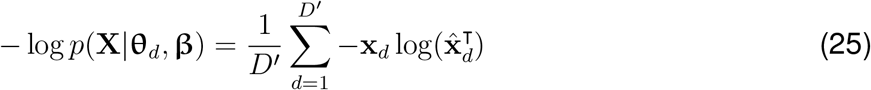

where *D*^*′*^ is the number of test individuals and 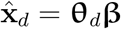

For medication imputation experiment, we masked the entire medication data of the test individuals and then reconstructing the medication matrix. This experiment mimics the scenario that we recommend relevant medications based only on individuals’ condition history.

In addition to reconstruction error, we also calculated recall and precision rates at the top 5 predictions: recall@5 and precision@5. Specifically, we sorted the predicted probabilities of medications and chose top 5 medications, of which we calculated recall and precision with respect to medications that the individual took as true labels. The recall and precision are calculated as: 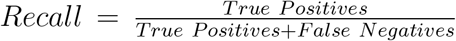 and 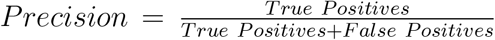, respectively.

#### CMK pain prediction

CMK pain was defined as musculoskeletal pain lasting at least three months localized to any of five body sites: knee, back, neck/shoulder, face or hip. The label information was obtained from the UKB pain questionnaire as described in **STAR Methods**.

We used individual topic mixture ***θ***_*d*_’s to predict whether a individual *d* has chronic musculoskeletal pain at the time of the visit as the binary outcome *y*_*d*_. We split data to 80% of training set and 20% of testing set. GETM was first trained with training set to get ***θ***^(*train*)^. Then we fit logistic regression model 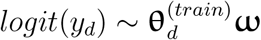, where ***ω*** is a *K ×* 1 vector of the linear coefficients corresponding to the predictive weights of the *K* topics.

To evaluate model performance, we first obtained ***θ***^(*test*)^ with testing set using the trained GETM (i.e., the encoder network parameters **W**_*θ*_) and then predicted CMK pain status using the trained logistic regression coefficients 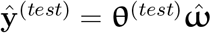.

#### Odds ratios of CMK-related conditions or medications

We calculated odds ratios of each condition and medication with respect to CMK pain as the outcome (**Supplementary Fig. ??**). The odds ratios were used for two purposes: (1) obtaining conditions and medications that are obvious for chronic musculoskeletal pain prediction and to be removed for some experiments; (2) constructing a baseline reference to construct a curated list of pain-related conditions and medication by the physician in our team. For each feature (condition or medication), we formed a contingency table as below. The odds ratios were calculated as 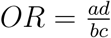.

| Feature \ Pain | Pain |  |
| --- | --- | --- |
|  | Yes | No |
| Yes | a | b |
| No | c | d |

#### Feature filtering schemes in removing obvious CMK-related conditions and medications

Let *𝒞* and *ℳ* be the complete sets of 443 conditions and 802 medications, respectively. Based on the odds ratios, we identified 50 conditions (*𝒞*1) and 150 medications (*ℳ*1) that are significantly associated with the CMK pain (**Supplementary Fig. ??**). A physician also provides a general pain related condition list (*𝒞*2), a musculoskeletal pain related condition list (*𝒞*3) and a general pain related medication set (*ℳ*2). We created three filtered condition sets: (1) *𝒞* − *𝒞*2; (2) *𝒞* − *𝒞*3; (3) *𝒞* − (*𝒞*1 ∪*𝒞*2), where using the setting notations *𝒞* − *𝒞*1 denotes condition set without *𝒞*1 and *𝒞*1 ∪ *𝒞*2 denotes the union of C1 and C2.

We also created two filtered medication sets: (1) *ℳ* − *ℳ*1; (2) *ℳ* − (*ℳ*1∪ *ℳ*2). These filtered condition sets and filtered medication sets formed six experiment settings plus the full set (**Supplementary Table ??**), which we used to explore CMK prediction performance as a function of reduced feature sets (**Fig**. 5).

#### Calculating importance of conditions and medications for CMK pain

Using the *K ×* 1 coefficients from logistic regression 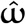 and the topic embeddings, we calculated the relevance of medications and conditions to CMK pain as follows (**Fig**. 6a,b):

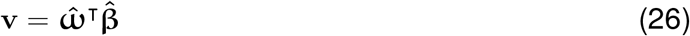

We selected the top *N ∈ {*10, 30, 50} medications and conditions from **v**^(*med*)^ and **v**^(*cond*)^. We then calculated the proportions of these top N medications or conditions overlapping with the pain-related lists created by a physician. The results are visualized as barplots in **Fig**. 6c and **Fig**. 6d.

#### Characterizing pain types from 7 body sites and pain all over the body

We used similar approaches as described for the CMK pain predictions to characterize other pain types. For all experiments, we used three different filtered feature sets of conditions and medications, which are (1) <u>all</u>: non-filtered conditions and medications, (2) <u>general</u>: physician-curated general pain-related conditions and medications were filtered, and (3) <u>least</u>: physician-curated / top odds ratio calculation conditions and medications were filtered.

For each pain type *p*, we calculated the importance weight that a medication or condition positively or negatively related to the pain using logistic regression coefficient for predicting that pain type 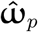 and GETM topic-feature mixture ***β*** of certain feature filtering regime (**Fig**. 6(b)). Specifically, the way to calculate importance weight *w*_*c,p*_ for a positive relation between a condition *c* and the pain type *p* is:

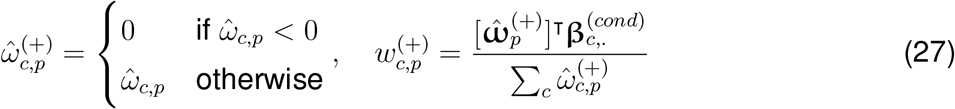

Similarly, we calculated importance weight for negative relationship between a condition *c* and pain type *p*:

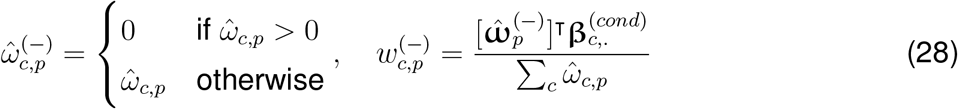

Calculation for the positive and negative relations between every medication and every pain type is the same.

### 10.3 QUANTIFICATION AND STATISTICAL ANALYSIS

Topic model were evaluated using topic quality scores as described in Section 8. The topic quality scores were computed for five repeated experiments with random initialization for each method (**Fig**. 3 and **Supplementary Fig. ??**). One-sided t-test were performed to compare the topic quality scores from the proposed Graph-embedded topic model (GETM) with the baseline methods. Prediction of chronic musculoskeletal pain phenotypes were evaluated using the receiver operating characteristic (ROC) curves and area under the ROC curve (AUROC) as well as precision-recall curves (PRC) and area under the PRC (AUPRC). Feature selection were performed using Fisher’s exact test as detailed above.

