## Supplementary Information for "A graph-embedded topic model enables characterization of diverse pain phenotypes among UK Biobank individuals"

### 1 Supplementary Figures

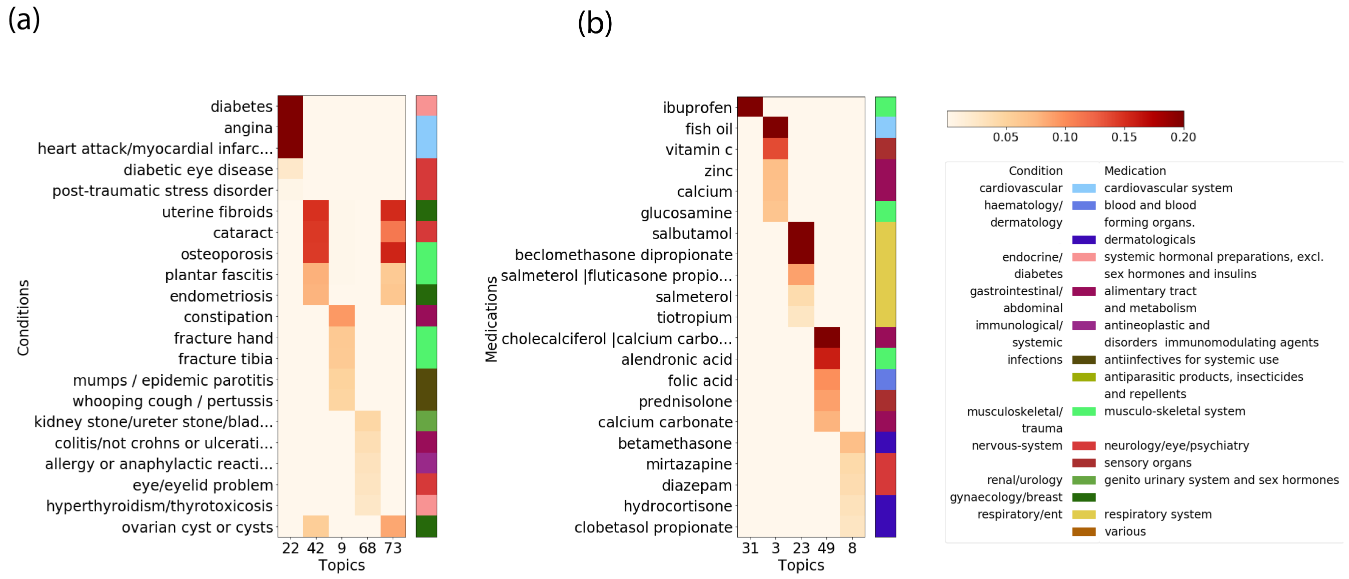


Figure S1: **Topic quality for ETM-condOnly and ETM-medOnly**. We chose 5 topics of diverse conditions and medications each for ETM-condOnly and ETM-medOnly models (**Supplementary Table** S1) which both used 75 topics. Then we visualized top 5 conditions or medications using heatmaps as in **Table** 2.

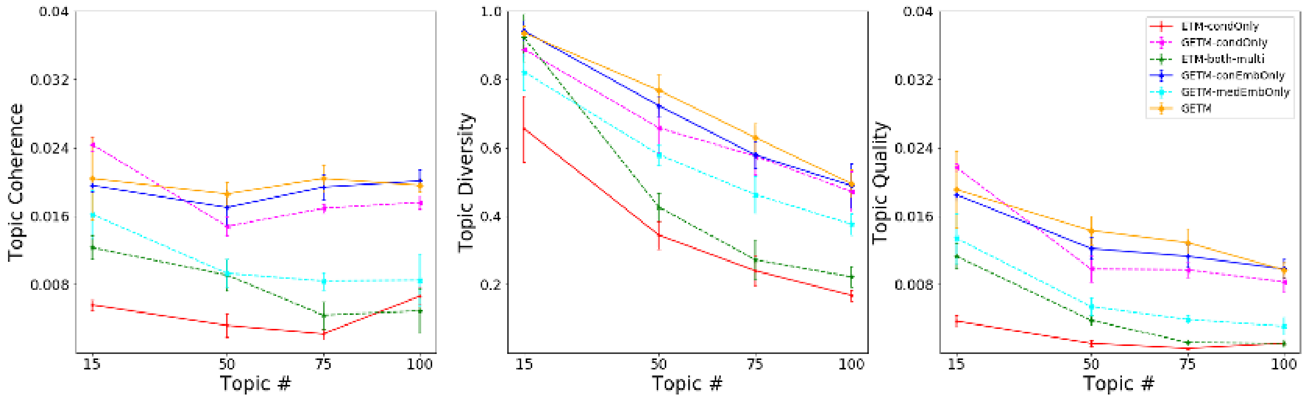


Figure S2: **Condition-specific topic quality evaluation.** We experimented with GTEM as well as 5 baseline models (**Supplementary Table** S1) with 4 predefined number of topics. To compute statistical significance between GETM and the baseline methods, we ran each model 5 times on the full UK Biobank data each with a different random initialization. The line plot displays the topic coherence, topic diversity, and topic quality, where the error bar indicates the standard deviations over the 5 experiments.

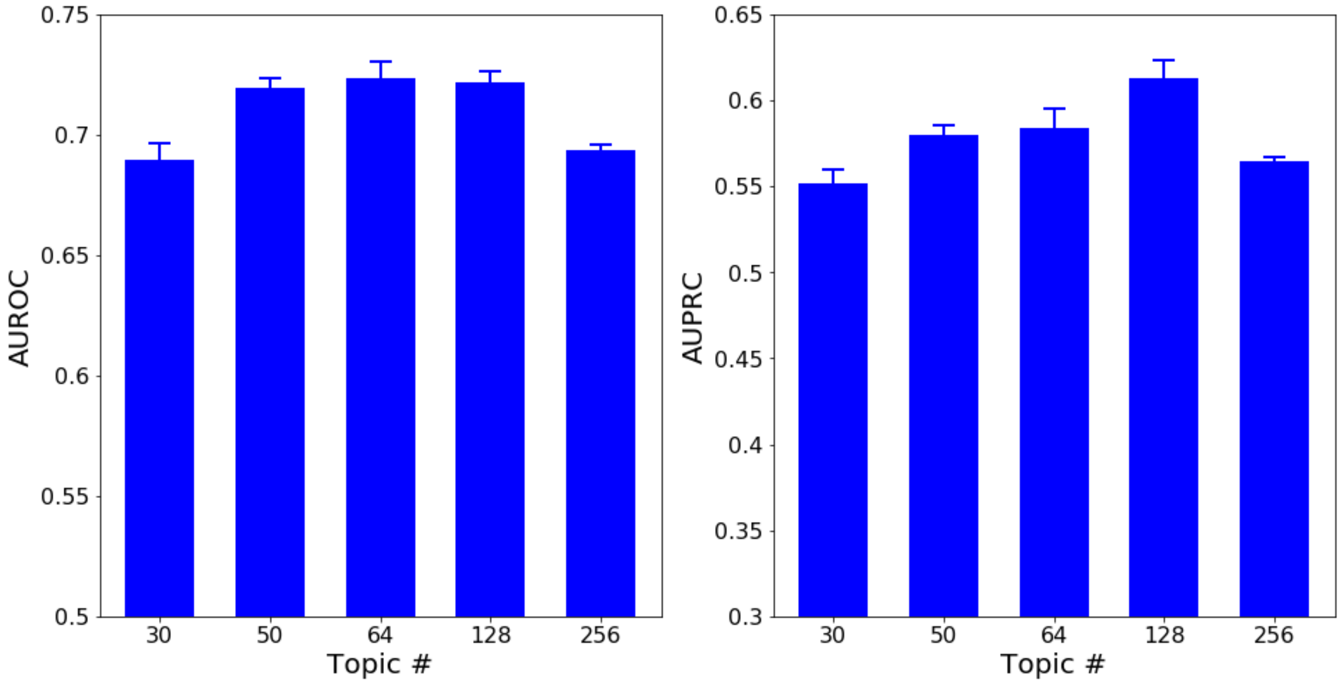


Figure S3: **Prediction performance of chronic musculoskeletal (CMK) pain using different topic numbers.** For each topic number K, we first trained a GETM on 80% of the training data without feature filtering (i.e., using 802 conditions and 443 medications to infer the K topic distributions). We then trained a logistic regression model using the infer patient topic mixture from the rained GETM as the input features to predict CMK pain labels on the 20% validation patients. We repeated the experiments 5 times to obtain the average AUROC and AUPRC and the standard deviations shown as the error bar.

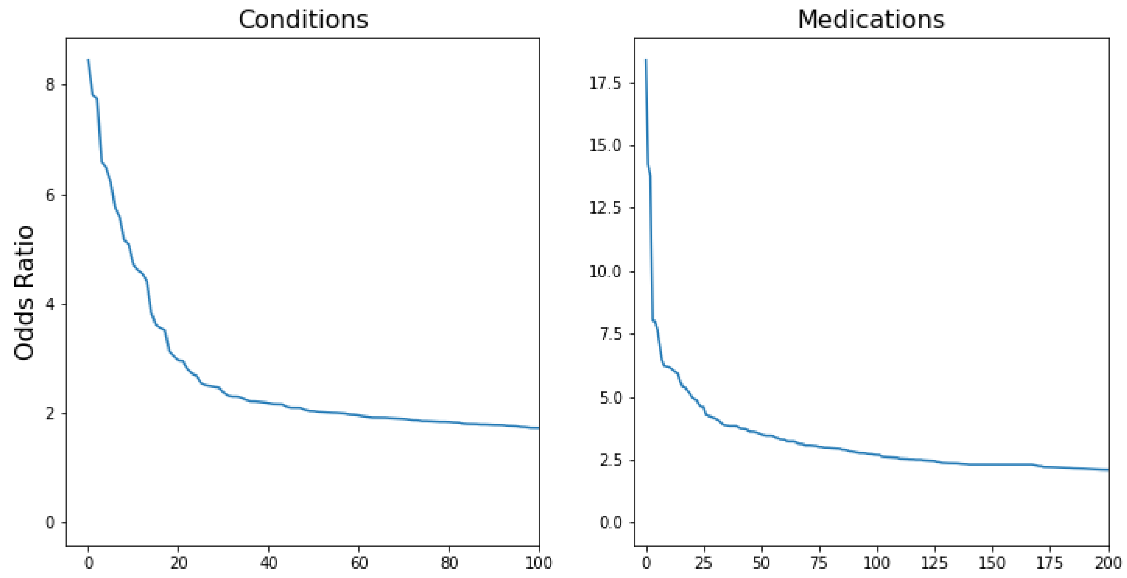


Figure S4: **Odds ratios for CMK pain**. The odds ratios of conditions and medications were calculated with respect to CMK pain. The odds ratios was used to remove some of the obvious conditions and medications in the prediction task illustrated in **Fig.** 5 in the main text.

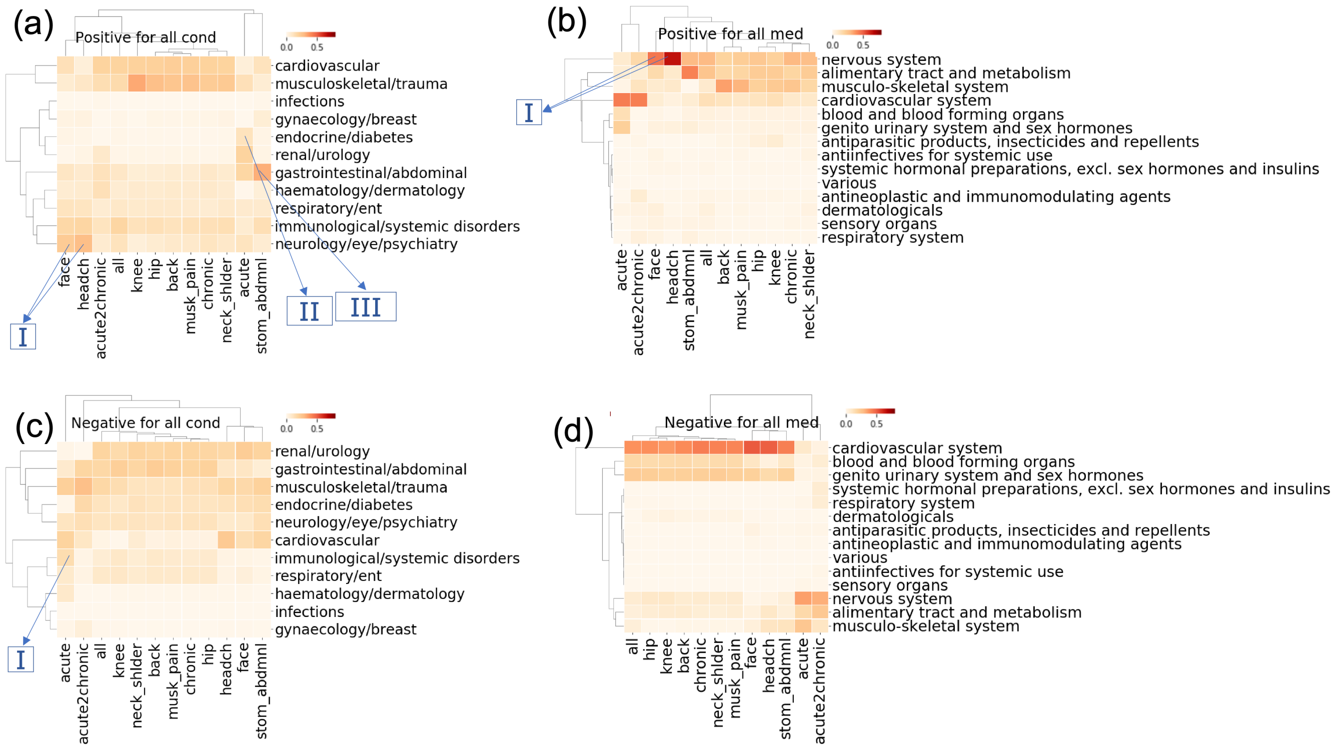


Figure S5: **Importance weights computed using non-filtered conditions and medications over all pain types**.For each pain type $p$, we calculated the importance weight that a medication or condition positively or negatively related to the pain using logistic regression coefficients for predicting that pain type $\hat{\omega}_p$ and GETM topic-feature mixture $\beta$ (**STAR Methods**). The physician-curated general pain-related conditions and medications were filtered.

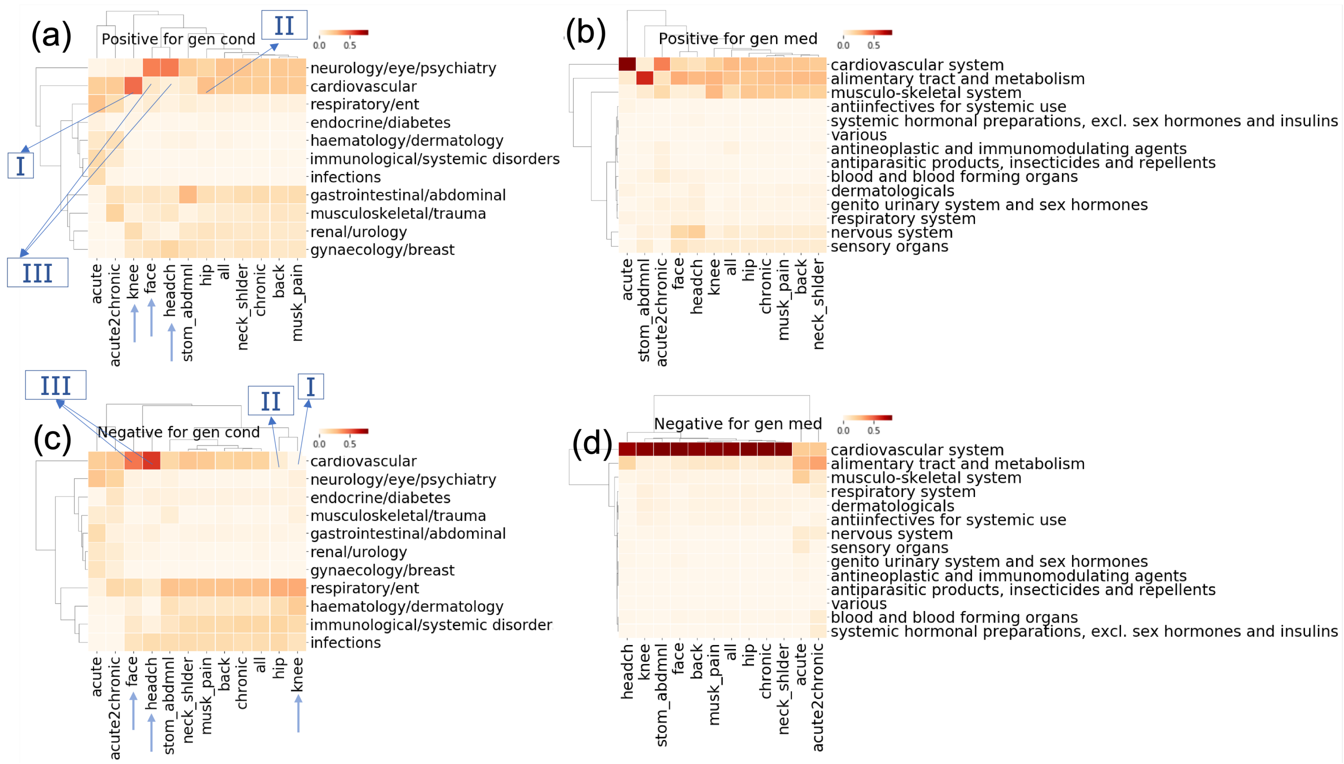


Figure S6: **Importance weights computed using physician-curated general pain-related filtered conditions and medications over all pain types**. For each pain type $p$, we calculated the importance weight that a medication or condition positively or negatively related to the pain using logistic regression coefficients for predicting that pain type $\hat{\omega}_p$ and GETM topic-feature mixture $\beta$ (**STAR Methods**).

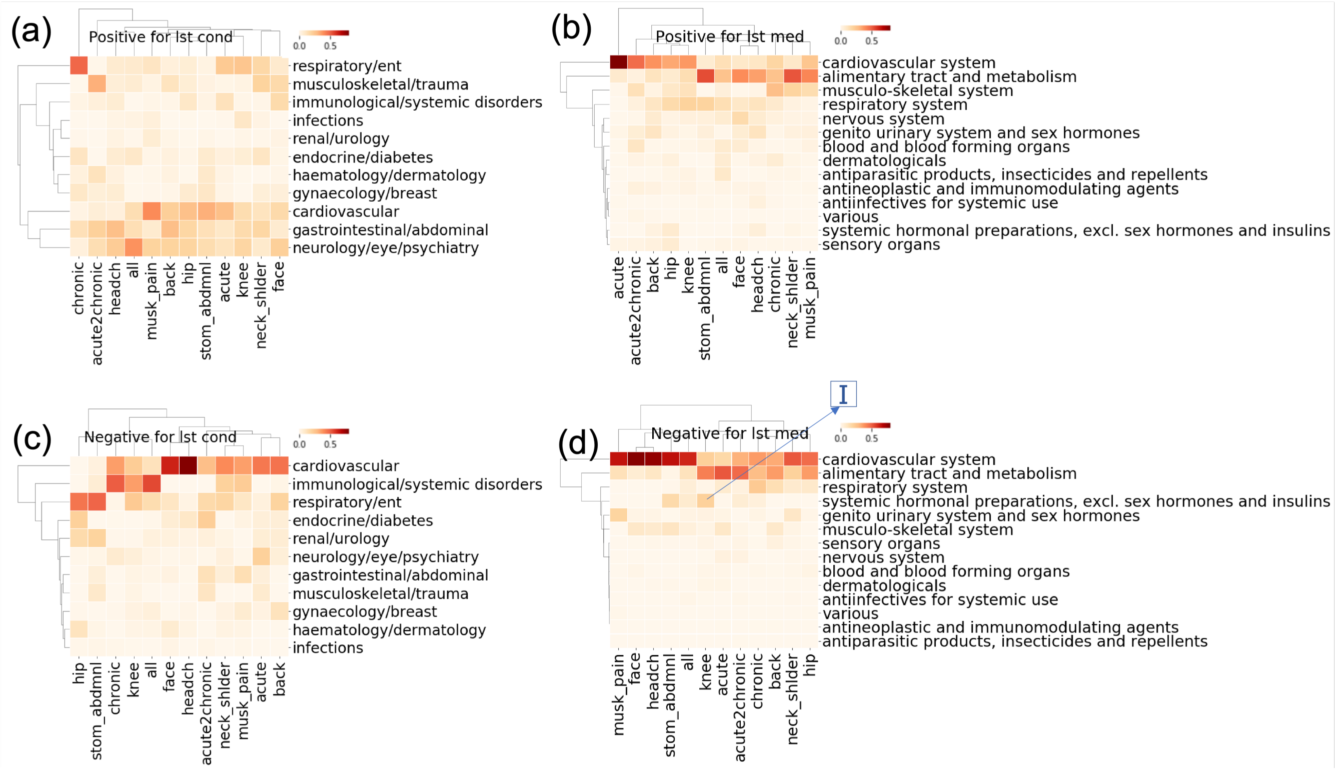


Figure S7: **Importance weights computed using physician-curated general pain-related filtered and top odds ratio calculated conditions and medications over all pain types**. For each pain type $p$, we calculated the importance weight that a medication or condition positively or negatively related to the pain using logistic regression coefficients for predicting that pain type $\hat{\omega}_p$ and GETM topic-feature mixture $\beta$ (**STAR Methods**).

### 2 Supplementary Tables

| Model Name | Model Description |
|---|---|
| ETM-condOnly | ETM modeling conditions only without node2vec pre-trained embedding |
| ETM-medOnly | ETM modeling medications only without pre-trained embedding |
| ETM-both-flat | Baseline ETM that takes both conditions and medications but treats them as the same type of features (i.e., single-view topics). |
| ETM-both-multi | ETM without using node2vec embedding but infers medication-specific and condition-specific topics (i.e., multi-view topics) |
| GETM-condOnly | GETM modeling only UKB conditions using the pre-trained node2vec embedding |
| GETM-medOnly | GETM modeling only UKB medications using the pretrained node2vec embedding |
| GETM-conEmbOnly | GETM using node2vec-embedding on condition but not on medications |
| GETM-medEmbOnly | GETM using node2vec-embedding on medication but not on conditions |
| GETM | The proposed full GETM model with node2vec-embeddings on both the conditions and medications |

Table S1: **The list of ablation models**. Abbreviation and description of the ablated models. All of the GETM models use the node2vec-embeddings pretrained on conditions or medications graphs. All of the ETM models do not use the node2vec-embeddings.

| Set # | Set Name | Set Description |
|---|---|---|
| 1 | m802c443 | containing all 802 medications; containing all 443 conditions |
| 2 | m680c389 | removing physician-curated general pain-related medications;<br>removing physician-curated musculoskeletal pain-related conditions |
| 3 | m680c351 | removing physician-curated general pain-related medications;<br>removing physician-curated general pain-related conditions |
| 4 | m680c322 | removing physician-curated general pain-related medications;<br>removing physician-curated general pain-related conditions<br>and top conditions correlated with chronic musculoskeletal pain from odds ratio calculation |
| 5 | m579c389 | removing physician-curated general pain-related medications<br>and top medications correlated with chronic musculoskeletal pain from odds ratio calculation;<br>removing physician-curated musculoskeletal pain-related conditions |
| 6 | m579c351 | removing physician-curated general pain-related medications<br>and top medications correlated with chronic musculoskeletal pain from odds ratio calculation;<br>removing physician-curated general pain-related conditions |
| 7 | m579c322 | removing physician-curated general pain-related medications<br>and top medications correlated with **chronic musculoskeletal pain** from odds ratio calculation;<br>removing physician-curated general pain-related conditions<br>and top conditions correlated with **chronic musculoskeletal pain** from odds ratio calculation |
| 8 | m666c340 | removing physician-curated general pain-related medications<br>and top medications correlated with **chronic neck/shoulder pain** from odds ratio calculation;<br>removing physician-curated general pain-related conditions<br>and top conditions correlated with **chronic neck/shoulder pain** from odds ratio calculation |
| 9 | m669c343 | removing physician-curated general pain-related medications<br>and top medications correlated with **chronic hip pain** from odds ratio calculation;<br>removing physician-curated general pain-related conditions<br>and top conditions correlated with **chronic hip pain** from odds ratio calculation |
| 10 | m674c341 | removing physician-curated general pain-related medications<br>and top medications correlated with **chronic back pain** from odds ratio calculation;<br>removing physician-curated general pain-related conditions<br>and top conditions correlated with **chronic back pain** from odds ratio calculation |
| 11 | m661c336 | removing physician-curated general pain-related medications<br>and top medications correlated with **chronic stomachache/abdominal pain** from odds ratio calculation;<br>removing physician-curated general pain-related conditions<br>and top conditions correlated with **chronic stomachache/abdominal pain** from odds ratio calculation |
| 12 | m671342 | removing physician-curated general pain-related medications<br>and top medications correlated with **chronic knee pain** from odds ratio calculation;<br>removing physician-curated general pain-related conditions<br>and top conditions correlated with **chronic knee pain** from odds ratio calculation |
| 13 | m670c337 | removing physician-curated general pain-related medications<br>and top medications correlated with **chronic headache** from odds ratio calculation;<br>removing physician-curated general pain-related conditions<br>and top conditions correlated with **chronic headache** from odds ratio calculation |
| 14 | m662c338 | removing physician-curated general pain-related medications<br>and top medications correlated with **chronic facial pain** from odds ratio calculation;<br>removing physician-curated general pain-related conditions<br>and top conditions correlated with **chronic facial pain** from odds ratio calculation |
| 15 | m665c348 | removing physician-curated general pain-related medications<br>and top medications correlated with chronic **all-body pain** from odds ratio calculation;<br>removing physician-curated general pain-related conditions<br>and top conditions correlated with chronic **all-body pain** from odds ratio calculation |
| 16 | m605c284 | removing physician-curated general pain-related medications<br>and top medications correlated with **chronic pain*** from odds ratio calculation;<br>removing physician-curated general pain-related conditions<br>and top conditions correlated with **chronic pain*** from odds ratio calculation |
| 17 | m640c332 | removing physician-curated general pain-related medications<br>and top medications correlated with **acute pain** from odds ratio calculation;<br>removing physician-curated general pain-related conditions<br>and top conditions correlated with **acute pain** from odds ratio calculation |

Table S2: List of runs conducted among UKB European descent study subjects including features from selected combinations of subsets of conditions (from a total of 443) and medications (from a total of 802).

| Label Name | Description |
|---|---|
| acute | acute pain on any of body sites |
| acute2chronic | acute pain on any of musculoskeletal body site at first visit and became chronic pain diagnosed at the following visits |
| chronic | chronic pain on any of body sites |
| back | chronic back pain |
| neck_shlder | chronic neck/shoulder pain |
| hip | chronic hip pain |
| knee | chronic knee pain |
| face | chronic facial pain |
| musk_pain | chronic musculoskeletal pain |
| headch | chronic headache |
| stom_abdmnl | chronic stomachache or abdominal pain |
| all | chronic pain on all over the body (all_over_body) |

Table S3: **Pain type label description**. The description for pain label name. Musculoskeletal pain sites includes face, knee, back, neck/shoulder and hip.

| Topic # / Algorithm | 15 | 50 | 75 | 100 | 15 | 50 | 75 | 100 | 15 | 50 | 75 | 100 |
|---|---|---|---|---|---|---|---|---|---|---|---|---|
| | Topic Coherence | | | | Topic Diversity | | | | Topic Quality | | | |
| ETM-medOnly | 0.2747 | 0.3302 | 0.3127 | 0.3292 | 0.8800 | 0.4370 | 0.2720 | 0.2640 | 0.2411 | 0.1450 | 0.0854 | 0.0872 |
| GETM-medOnly | 0.4827 | 0.6720 | 0.7131 | 0.7072 | 0.9680 | 0.8310 | **0.6940** | 0.5720 | 0.4667 | 0.5585 | **0.4951** | 0.4045 |
| ETM-both-multi | 0.3693 | 0.3600 | 0.3661 | 0.3612 | 0.8460 | 0.4990 | 0.2880 | 0.2160 | 0.3135 | 0.1796 | 0.1051 | 0.0779 |
| GETM-conEmbdOnly | 0.3760 | 0.4532 | 0.3366 | 0.4012 | 0.9660 | 0.5610 | 0.4560 | 0.3680 | 0.3629 | 0.2539 | 0.1536 | 0.1472 |
| GETM-medEmbOnly | 0.4827 | 0.6912 | **0.7739** | **0.7812** | **0.9840** | 0.7720 | 0.600 | 0.484 | 0.4747 | 0.5339 | 0.4645 | 0.3780 |
| GETM | **0.5289** | **0.7012** | 0.7104 | 0.7236 | 0.9680 | **0.8640** | 0.6800 | **0.5600** | **0.5109** | **0.6061** | 0.4823 | **0.4050** |

Table S4: **Medication-specific topic evaluation**. We have calculated topic coherence, topic diversity and topic quality in terms of medications for 6 algorithms using different topic numbers (model details in **Supplementary Table** S1. Condition-specific topic evaluation is in **Supplementary Table** S5.

| Topic # / Algorithm | 15 | 50 | 75 | 100 | 15 | 50 | 75 | 100 | 15 | 50 | 75 | 100 |
|---|---|---|---|---|---|---|---|---|---|---|---|---|
| | Topic Coherence | | | | Topic Diversity | | | | Topic Quality | | | |
| ETM-condOnly | 0.0056 | 0.0032 | 0.0022 | 0.0066 | 0.6560 | 0.3440 | 0.2400 | 0.1680 | 0.0037 | 0.0011 | 0.0005 | 0.0011 |
| GETM-condOnly | **0.0244** | 0.0148 | 0.0169 | 0.0176 | 0.8880 | 0.6580 | 0.5760 | 0.4720 | **0.0217** | 0.0098 | 0.0097 | 0.0083 |
| ETM-both-multi | 0.0123 | 0.0091 | 0.0044 | 0.0049 | 0.9220 | 0.4260 | 0.2720 | 0.2220 | 0.0113 | 0.0038 | 0.0012 | 0.0011 |
| GETM-conEmbOnly | 0.0196 | 0.0170 | 0.0194 | **0.0201** | 0.9420 | 0.7230 | 0.5790 | 0.4890 | 0.0185 | 0.0122 | 0.0113 | 0.0098 |
| GETM-medEmbOnly | 0.0162 | 0.0093 | 0.0084 | 0.0085 | 0.8220 | 0.5800 | 0.4630 | 0.3760 | 0.0134 | 0.0054 | 0.0039 | 0.0031 |
| GETM | 0.0204 | **0.0186** | **0.0204** | 0.0196 | **0.9360** | **0.7680** | **0.6300** | **0.4950** | 0.0191 | **0.0143** | **0.0129** | **0.0097** |

Table S5: **Condition-defined topic quality**. We have calculated topic coherence, topic diversity and topic quality in terms of conditions for 6 algorithms using different topic numbers. The description of algorithm name is in **Supplementary Table** S1.

| Topic # / Algorithm | 15 | 50 | 75 | 100 |
|---|---|---|---|---|
| | Reconstruction Error | | | |
| ETM-medOnly | 8.54 | 8.24 | 8.10 | 7.56 |
| GETM-medOnly | 7.84 | 7.46 | 6.13 | 6.99 |
| ETM-both-multi | 7.35 | 7.40 | 7.63 | 6.66 |
| GETM-condEmbOnly | 6.70 | 6.85 | 6.83 | 6.74 |
| GETM-medEmbOnly | 7.71 | 7.02 | 6.98 | 6.28 |
| GETM | **6.34** | **5.70** | **5.69** | **5.53** |
| Proportion | 12.23 | | | |

Table S6: **Imputation error of masked medications**. The 50% of test data was randomly masked. Then we reconstructed the matrix with learned $\theta, \alpha$ and $\rho$. The reconstruction error (i.e. negative log-likelihood) was calculated for the held-out data. Same medication data was used for all algorithms. Same models and same data were used as in **Table** 2 for ETM-both-multi, GETM-condEmbOnlym GETM-medEmbOnly and GETM. Description of algorithm names are in **Supplementary Table** S1. As another baseline method, we evaluated the performance of filling in masked conditions based on their overall proportion over all of the UKB population.

| Column 1 | Column 2 |
|---|---|
| evening primrose oil | multivitamins minerals and combinations |
| antiadrenergic agents | opioid - non-opioid combinations |
| dopaminergic agents | oral hypoglycemics |
| insulins and analogues | anxiolytics hypnotics and sedatives |
| antifungals | vasodilators |
| antispasmodics | decongestants and antiallergics |
| antivertigo preparations | chondroitin |
| topical products for joint and muscular pain | cardiac glycosides |
| laxatives emollients and protectives | hormones and related agents |
| antimigraine preparations | antiepileptics |
| beta blocking agents | fish oil |
| diuretics | antipsoriatics |
| other topical preparations | nsaid |
| parasympathomimetics | ginkgo biloba |
| anticholinergic agents | calcium |
| glucosamine | paracetamol |
| calcium channel blockers | antidepressants |
| bisphosphonates | all other therapeutic products |
| iron and other antianemic preparations | antigout preparations |
| thyroid preparations | antacids drugs for peptic ulcer and gerd |
| angiotensin ii receptor blockers | corticosteroids |
| antithrombotic agents | other psychotherapeutic agents |
| antiarrhythmics - i and iii | antimalarials |
| intestinal antiinflammatory agents | antibacterials |
| estrogens progestogens and combinations | ace inhibitors |
| immunosuppressants | antivirals |
| urologicals | opioids |
| lipid modifying agents | salicylic acid and derivatives |
| ophthalmologicals | adrenergic agents |
| antihistamines | |

Table S7: **External medication categories** All 59 external medication categories are listed.
